# Is FAM19A5 an adipokine? Peripheral FAM19A5 in wild-type, FAM19A5 knockout, and LacZ knockin mice

**DOI:** 10.1101/2020.02.19.955351

**Authors:** Hoyun Kwak, Eun-Ho Cho, Eun Bee Cho, Yoo-Na Lee, Anu Shahapal, Hyo Jeong Yong, Arfaxad Reyes-Alcaraz, Yongwoo Jeong, Yerim Lee, Minhyeok Lee, Nui Ha, Sitaek Oh, Jae Keun Lee, Won Suk Lee, Won Kyum Kim, Jong-Ik Hwang, Jae Young Seong

## Abstract

FAM19A5 is a novel secretory protein primarily expressed in the brain. However, a recent study reported that FAM19A5 is an adipocyte-derived adipokine that regulates vascular smooth muscle function through sphingosine-1-phosphate receptor 2 (S1PR2). In our study, we investigated FAM19A5 transcript and protein levels in peripheral tissues, including adipose tissues, from wild-type, FAM19A5 knockout, and FAM19A5 LacZ knockin mice. We found that FAM19A5 transcript levels in the central nervous system were much greater than those in any of the peripheral tissues, including adipose tissues. Furthermore, the FAM19A5 protein levels in adipose and reproductive tissues were below detectable limits for Western blot analysis. Additionally, we found that the FAM19A5 protein did not interact with S1PR2 in terms of G protein-mediated signal transduction, β-arrestin recruitment, or ligand-mediated internalization. Taken together, our findings revealed basal levels of FAM19A5 transcripts and proteins in peripheral tissues, confirming its primary expression in the central nervous system and lack of significant interaction with S1PR2.

## INTRODUCTION

FAM19A5 is a member of the FAM19A protein family (also called the TAFA family) and is highly expressed in the brain (Tom Tang et al., 2004). The FAM19A5 gene encodes a secretory protein with a high degree of amino acid sequence identity across vertebrate species (Jeong et al., 2021), suggesting its functional relevance. Transcripts for FAM19A5 are abundant in the brain and relatively rare in peripheral tissues (Tom Tang et al., 2004). A recent study using FAM19A5-LacZ knockin (KI) mice revealed the expression patterns of FAM19A5 during embryogenesis and in the adult brain (Shahapal et al., 2019). FAM19A5 is expressed in the ventricular zone and ganglionic eminence during early brain development, suggesting a role in the proliferation and differentiation of neural stem cells and oligodendrocyte precursor cells (OPCs) (Shahapal et al., 2019). In the adult brain, X-gal staining combined with immunostaining for cell-type markers revealed that FAM19A5 is expressed in diverse cell types, including neurons, astrocytes, and OPCs (Shahapal et al., 2019). These findings are consistent with single-cell RNA sequencing data from mouse and human brains (Zeisel et al., 2018; Karlsson et al., 2021; Sjöstedt et al., 2020; Mathy et al., 2019).

The results from FAM19A5 knockout (KO) and FAM19A5 LacZ KI mouse models suggest that FAM19A5 may affect the formation and elimination of synapses in neurons (Huang et al., 2021; Shahapal et al., 2024). Furthermore, we recently demonstrated that FAM19A5 contributes to synapse elimination by binding to LRRC4B, a postsynaptic cell adhesion molecule, through the FAM19A5 binding domain within LRRC4B (Kim et al., 2023). LRRC4B is known to participate in synapse formation by binding to PTPRF, a cell adhesion molecule in the presynapse (Woo et al., 2009). However, when FAM19A5 binds to LRRC4B, the interaction between LRRC4B and PTPRF is inhibited. Conversely, treatment with NS101, a monoclonal antibody that inhibits the function of FAM19A5, restored the lost synapses. This action of NS101 also restored the loss of synapses and cognitive function in mouse models of Alzheimer’s disease (Kim et al., 2023).

In addition to its functional importance in the brain, a recent study showed that FAM19A5 suppressed neointima formation in injured rat carotid arteries by interacting with sphingosine-1-phosphate receptor 2 (S1PR2) (Wang et al., 2018). This study indicated that FAM19A5 is primarily produced in adipose tissue. qRT□PCR analysis of human tissue samples revealed that the FAM19A5 mRNA level in adipose tissue was greater than that in brain tissue. Furthermore, immunohistochemistry and ELISA for FAM19A5 have shown high levels of the FAM19A5 protein in human and rodent adipose tissues (Wang et al., 2018). Additionally, FAM19A5 mRNA and protein levels are significantly downregulated in obese and diabetic animals (Wang et al 2018), suggesting a causal relationship between decreased FAM19A5 levels and metabolic disease conditions. However, this result partially contradicts recent public database reports on human tissue RNA expression, which show that FAM19A5 is expressed in female and male reproductive tissues, including the ovaries, uterus, and testis, although its absolute levels in these tissues are significantly lower than those in brain tissue (Karlsson et al., 2021; Sjöstedt et al., 2020).

In the present study, we examined the basal levels of FAM19A5 transcripts and proteins in a variety of peripheral tissues via qRT□PCR and Western blotting. The methods were verified with FAM19A5 KO mice. We also identified the cell types that express FAM19A5 via X-gal staining of peripheral tissues from FAM19A5-LacZ KI mice, as previously described (Shahapal et al., 2019). Contrary to a previous report (Wang et al., 2018), this study also demonstrated that FAM19A5 transcript and protein levels in adipose tissues are very low compared to those in brain tissues and that S1PR2 does not interact with the FAM19A5 protein, as determined by receptor-mediated signal transduction, β-arrestin recruitment, and ligand-mediated receptor internalization.

## MATERIALS AND METHODS

### Animals

C57BL/6J mice were purchased from Nara Biotech (Korea) or Orient Bio, Inc. (Korea). Mice were housed under temperature-controlled (22–23°C) conditions with a 12-h light/12-h dark cycle. The mice were given standard chow and water ad libitum. All animal experiments were designed to use the fewest mice possible, and anesthesia was administered when necessary. All animal procedures were approved by the Institutional Animal Care and Use Committee of Korea University (KOREA-2019-0076 and KOREA-2019-0032).

### Generation of FAM19A5 knockout (KO) mice

FAM19A5 KO mice were generated by ToolGen, Inc. (Korea) using the CRISPR/Cas9 system (Lee et al., 2018). Briefly, single-guide RNAs (sgRNAs) were designed to target a 5’-UTR sequence of exon 1 (left sgRNA, ccgtctctgtcgccatccagagg E1-1) and the 3’-UTR sequence of exon 4 (right sgRNA, cttggcacttaactcccagatgg). Cas9 mRNA and sgRNAs were microinjected into the cytoplasm of C57BL/6J mouse zygotes, and the resulting embryos were transferred into the oviducts of IcrTac:ICR pseudopregnant foster mothers to produce live founder mice. Founder mice with mutant alleles were screened using genomic DNA PCR to amplify the genomic region spanning the sgRNA target sites. The primer sequences for the founder screening PCR were as follows: forward, GGGGGTCCCAAGTCACCTAAC; reverse, AAGAACTTGGGAGACAGGCAAA. The PCR products from the founder mice were cloned, and the corresponding mutations were identified by direct-sequencing analysis (Bionics Co., Ltd., Korea). Founder mice with a mutant allele that lacked the 125,000 bp sequence containing exons 1 to 4 were bred with wild-type mice. Germline transmission of the mutant allele was determined by genotyping with the following primers: gF1, TCGGTTCACTTTCCGGATCAAT; gR1, AAGAACTTGGGAGACAGGCAAA; and gF2, TCCTGGGAGAGGGGAATAGTTT. Homozygous FAM19A5 KO mice were generated by heterozygous intercross breeding.

### FAM19A5-LacZ-KI mice

*FAM19A5-LacZ* KI mice were generated by the UC Davis Mouse Biology Program as previously described (Shahapal et al., 2019). The gene-trap method using *LacZ* as a reporter gene was employed to visualize FAM19A5 expression in tissue sections (Mountford et al., 1994). Briefly, the target vector containing the IRES-*lacZ* gene was inserted in front of exon 4 of the *FAM19A5* gene. The *LacZ* gene is expressed independently of the target *FAM19A5* gene due to an IRES element. This *FAM19A5*-targeting vector was delivered to embryonic stem cells by electroporation. We confirmed the incorporation of the vector into the target chromosome by genotyping and chromosome counting of transgenic embryonic stem cells. Selected transgenic embryonic stem cells were injected into blastocysts, and the embryos were implanted into the uterus of female recipient mice. We performed a germline transmission test to check for stable germline expression in the chimeric generation. The generated *FAM19A5-LacZ* KI chimeric mice were backcrossed onto a C57BL/6J genetic background.

### Quantitative real□time polymerase chain reaction (qRT□PCR) analysis

TRIzol reagent (Invitrogen) was used to isolate total RNA from mouse tissues. The analyzed tissues included the cerebral cortex, hippocampus, spinal cord, heart, aorta, kidney, lung, stomach, small intestine, large intestine, pancreas, liver, salivary gland, thymus, spleen, bone marrow, adrenal gland, pituitary gland, thyroid gland, skeletal muscle, white adipose tissue, brown adipose tissue, and skin. One microgram of RNA was reverse□transcribed into complementary DNA with a RevertAid First Strand cDNA Synthesis Kit (Thermo Scientific). The sequences of primers used for qRT□PCR were as follows: mFAM19A5-Iso1-F, GTCCTCAACTTTTTGGGCATTC; mFAM19A5-Iso2-F, GCGATGCAGCTCCTGAAG; mFAM19A5-Iso1,2-R, CCCGGTCTAGGGTCACAA; mFAM19A5-Total-F, GGCAGATAGCAGGCACCACT; mFAM19A5-Total-R, GCTGCGATTGTCAGGAGACC; mGAPDH-F, ATCCTG CACCACCAACTGCT; and mGAPDH-R, GGGCCATCCACAGTCTTCTG. The CFX96 Touch RT□PCR detection system using SsoAdvanced Universal SYBR Green Supermix (Bio-Rad) was used for qRT□PCR. Gene expression was normalized to GAPDH levels, and the relative quantity of mRNA was calculated using the comparative Cq method. Possible genomic DNA contamination for all tissues was assessed by performing identical PCRs on samples that had not been reverse transcribed and showed either no contamination or negligible levels of contamination that did not interfere with qRT□PCR.

### X-gal staining

X-gal staining was performed as previously described (Shahapal et al., 2019) using 10-week-old adult wild-type (WT) littermates (2 males) and *FAM19A5-LacZ* KI heterozygote (1 male) and homozygote (3 males and 1 female) mice. Mice were perfused with normal saline and 4% paraformaldehyde in 1× phosphate-buffered saline (PBS). Subsequently, brain and peripheral tissues were isolated, fixed in 4% paraformaldehyde for 3 h at 4°C and cryoprotected with 30% sucrose in 1× PBS for 36 h at 4°C. The cryoprotected tissues were embedded in optimal compound temperature (OCT) solution and 30% sucrose (2:3 ratio). The embedded blocks were sectioned at 20 μm using a cryostat (Leica). Cryosections that were mounted on glass slides were washed twice with 1× PBS for 5 min each, permeabilized with 0.01% sodium deoxycholate and 0.02% Igepal CA-630 in 1× PBS for 15 min and incubated in an X-gal solution for 24 h at 37°C in the dark. The X-gal solution contained 1 mg/ml X-gal, 2 mM magnesium chloride, 5 mM ethylene glycol-bis(2-aminoethylether)-N,N,N′,N′-tetraacetic acid, 5 mM potassium ferrocyanide, 5 mM potassium ferricyanide, 0.01% sodium deoxycholate, and 0.02% Igepal CA-630 in 0.1 M phosphate buffer (PB) at pH 7.4. X-gal-stained sections were washed twice with 1X PBS for 5 min each, counterstained with Mayer’s hematoxylin, dehydrated with graded ethanol and xylene, and mounted with a cover glass.

The tissue-specific X-gal signal was scored according to the following criterion: no X-gal signal in either WT or *FAM19A5-LacZ*-KI mice was indicated by “-”. The nonspecific X-gal signal detected in both the WT and *FAM19A5-LacZ*-KI mice, which was likely due to endogenous β-gal, was represented by “Δ”. An ambiguous signal in both WT and *FAM19A5-LacZ* KI mice was indicated by “Δ/+”. Ambiguity was defined as a signal in *FAM19A5-LacZ* KI mice that appeared stronger than the signal in WT mice or a negligible signal that was found only in *FAM19A5-LacZ* KI mice. A positive signal observed only in *FAM19A5-LacZ* KI mice but not in WT mice is denoted by “+”. Among the samples with a positive signal, “1+” indicated a faint positive signal with a sparse, cellular blue precipitate, “2+” indicated a weak positive signal with one or more blue precipitate areas per cell, “3+” indicated a moderate positive signal that was observed in many cells, and “>4+” indicated a robust positive signal.

### Purification and fluorescence labeling of the FAM19A5 protein

Recombinant N-terminal His-tagged FAM19A5 (N-HIS-FAM19A5) with a tobacco etch virus protease recognition sequence was cloned and inserted into the vector pCAG1.1 and expressed in Expi293F cells (Gibco). The supernatant of the transfected cells encoding N-HIS-FAM19A5 was collected for Ni-NTA affinity chromatography (Cytiva). Purified N-HIS-FAM19A5 was then digested overnight at 30°C with AcTEV protease (Invitrogen) to remove the His-tag. Following the cleavage reaction, the digestion products were verified by SDS□PAGE and Western blot analysis. Purified FAM19A5 was used as a Western blotting control. Next, the purified FAM19A5 was labeled with Cy3 using a protein labeling kit (Abcam) according to the manufacturer’s instructions. Following the reaction, the FAM19A5-Cy3 conjugates were buffer-exchanged into phosphate-buffered saline (PBS) and concentrated using Amicon filters (Millipore).

### Generation of anti-FAM19A5 antibodies

To generate anti-FAM195 chimeric human/chicken monoclonal antibodies, chickens (*Gallus gallus* domesticus) were immunized with purified N-terminal His-tagged FAM19A5 protein. Total RNA was extracted from the spleen, bursa of Fabricius, and bone marrow of the immunized chickens, and cDNA was synthesized. From the synthesized cDNA, a single-chain variable fragment (scFv) library was constructed with the pComb3X-SS vector system (Scripps Research Institute, USA). The helper phage VCM13 was added to the culture media for the extraction of phagemid DNA and phage-displayed scFv. Biopanning was performed using recombinant N-HIS-FAM19A5-coated magnetic beads with the phage-displayed scFv. Phages bound by biopanning were amplified and used for a second round of biopanning. Biopanning was repeated five times, and each time, the number of washes was increased to identify the phage with the highest affinity. ELISA was used to test the affinity of randomly selected phage-displayed scFv clones from biopanning for recombinant FAM19A5. The human C_К_ gene was linked to the anti-FAM19A5 antibody sequence that encodes the light chain variable region, and the CH1, CH2, and CH3 genes of the human immunoglobulin isotype IgG1 were added to the heavy chain variable region (GenScript). Vectors that encoded the anti-FAM19A5-IgG1 antibody were transfected into HEK293F cells, and the recombinant antibody was purified using Protein A beads (RepliGen). Anti-FAM19A5 antibodies that recognized the epitopes formed at the N-terminal and C-terminal regions were generated and called N-A5-Ab and C-A5-Ab, respectively. The purified antibodies were conjugated with HRP using an antibody labeling kit (Abcam) according to the manufacturer’s instructions.

### Western blot analysis

Mouse tissue samples were homogenized in buffer containing 20 mM Tris-HCl (pH 7.5), 500 mM NaCl, 5 mM MgCl_2_, and protease inhibitor cocktail (Thermo Scientific). The cell lysates were prepared in lysis buffer containing 20 mM Tris-HCl (pH 7.5), 150 mM NaCl, 0.5% NP-40, and protease inhibitor cocktail. Protein quantification of the homogenized tissues and cell lysates was performed using the Pierce BCA Protein Assay (Thermo Scientific). Brain homogenates, cell lysates, and cell culture media were then resolved on Tris-glycine gels (Invitrogen) and transferred to prewetted PVDF blotting membranes using a Trans-Blot Turbo apparatus (Bio-Rad). The membrane was blocked in Tris-buffered saline containing 0.05% Tween 20 and 5% skim milk, followed by overnight incubation with HRP-conjugated N-A5-Ab at 4°C. The blots were washed three times with Tris-buffered saline (TBS) containing 0.05% Tween 20. After the application of ECL reagents (Thermo Scientific), the immunoreactive bands were visualized (photographed) using a Mini HD9 (UVerytec) system.

### ELISA

To measure FAM19A5 in the CSF and plasma of WT and FAM19A5 KO mice, 96-well microplates were coated with 6xHis-TEV-LRRC4B (453-576) protein (1 µg/ml in 50 mM carbonate buffer, pH 9.6) and incubated overnight at 4°C. The next day, the plates were washed twice with 300 µL of washing buffer (PBS with 0.05% Tween 20) using a microplate washer (Tecan) and tapped dry. Then, 200 μL of blocking buffer was added to each well, and the plates were incubated at 37°C for 1 hour. After two additional washes, 100 μL of standard solution and sample were added to each well and incubated at room temperature for 1 hour. Following five washes, 100 μL of HRP-conjugated C-A5-Ab (0.2 μg/ml in blocking buffer) was added to each well and incubated at 37°C for 1 hour. Then, 100 μL of TMB solution (Thermo Scientific) was added to each well and incubated at room temperature for 20 minutes. The reaction was stopped with 100 μL of 1 N sulfuric acid, and the OD was measured at 450 nm using a microplate reader (Molecular Devices).

### SRE-luciferase assay for S1PR2 activation

The SRE-luciferase (SRE-luc) vector containing a single copy of the serum response element (SRE: CCATATTAGG) conjugated to luciferase was purchased from Stratagene. The cDNAs for human S1PR2 were obtained from BRN Science, Inc. The cDNA genes were inserted into the *EcoRI* and *XhoI* sites of the pcDNA3.1 vector (Invitrogen). Previously, we constructed a HEK293 cell line that stably expressed G_qi_ and allowed for the induction of G_q_-dependent signaling pathways upon activation of a G_i_-coupled receptor (Conklin et al., 1993; Kim et al., 2014). For the luciferase assays, HEK293G_qi_ cells were seeded on 48-well plates. Transfections were performed using a mixture of the SRE-luc reporter construct, S1PR2 expression plasmid, and Lipofectamine 2000 (Invitrogen), and the cells were incubated in Opti-MEM (Gibco) for 20 min before being applied to the cells. After transfection, the cells were incubated in serum-free DMEM for 16∼18 h and then treated with either sphingosine-1-phosphate (S1P; Sigma□Aldrich) or purified FAM19A5 protein for 6 h. The cells were lysed by adding 100 μl of lysis buffer to the wells. Luciferase activity was determined by analyzing 50 μl of each cell extract in a luciferase assay system according to the standard protocol for the Synergy 2 Multi-Mode Microplate Reader (BioTek).

### Confocal imaging of S1PR2 and FAM19A5 internalization

HEK293 cells were seeded at a density of 4.5 × 10^4^ cells/well on poly-L-lysine-coated plastic film coverslips in 12-well plates. The following day, the cells were transfected with S1PR2-GFP plasmids. One day posttransfection, the cells were incubated in serum-free MEM for 16 h prior to treatment with S1P (Sigma□Aldrich) and FAM19A5. Cell images were taken 30 min after ligand treatment using a confocal laser scanning microscope (Leica TCS-SP8).

### NanoLuc luciferase complementation assay

The NanoLuc luciferase complementation assay to detect S1PR2 and β-arrestin interactions was performed as previously described (Reyes-Alcaraz et al., 2018; Reyes-Alcaraz et al., 2019). Briefly, the NanoBit starter kit containing the plasmids and the necessary reagents to complete the structural complementation assays in this study were a gift from the Promega Company. HEK293 cells were maintained in DMEM supplemented with 10% FBS and penicillin□streptomycin (Gibco). One day prior to transfection, the cells were seeded in poly-L-lysine-coated 96-well plates at a density of 2.5 × 10^4^ cells per well. A mixture containing 100 ng of β-arrestin construct with SmBit, 100 ng of S1PR2 with LgBit or Nluc, and 0.4 μl of Lipofectamine 2000 was prepared and added to each well. Six hours posttransfection, the medium was aspirated and replaced with serum-free DMEM. Twenty-four hours posttransfection, the medium was aspirated and replaced with 100 μl of Opti-MEM at room temperature. After a 10 min incubation at room temperature, 25 μl of furimazine substrate was added, and luminescence measurements were recorded every minute for 10 min. Next, 10 μl of ligand was added to each well, and luminescence measurements were recorded immediately for 30 min at 1 min intervals (Synergy H1 Hybrid Multi-Mode Reader, BioTek).

## RESULTS

### FAM19A5 mRNA levels in peripheral and brain tissues

The human and mouse FAM19A5 genes can produce two transcript isoforms, isoform 1 and isoform 2, due to the alternative use of the first exon, exon 1-1 and exons 1-2 (Wang et al 2018). To determine the FAM19A5 transcript levels, tissue samples from the central and peripheral nervous systems were analyzed using qRT□PCR. Additionally, tissue samples from other peripheral tissues were also analyzed by qRT□PCR. Transcript levels were detected with specific primer sets for FAM19A5 isoform 1, FAM19A5 isoform 2, and the total transcript (Fig. 1A). The total FAM19A5 transcript levels were approximately equal to the sum of the isoform 1 and isoform 2 levels (Fig. 1B, C, D, and Table 1).

**Fig. 1.**
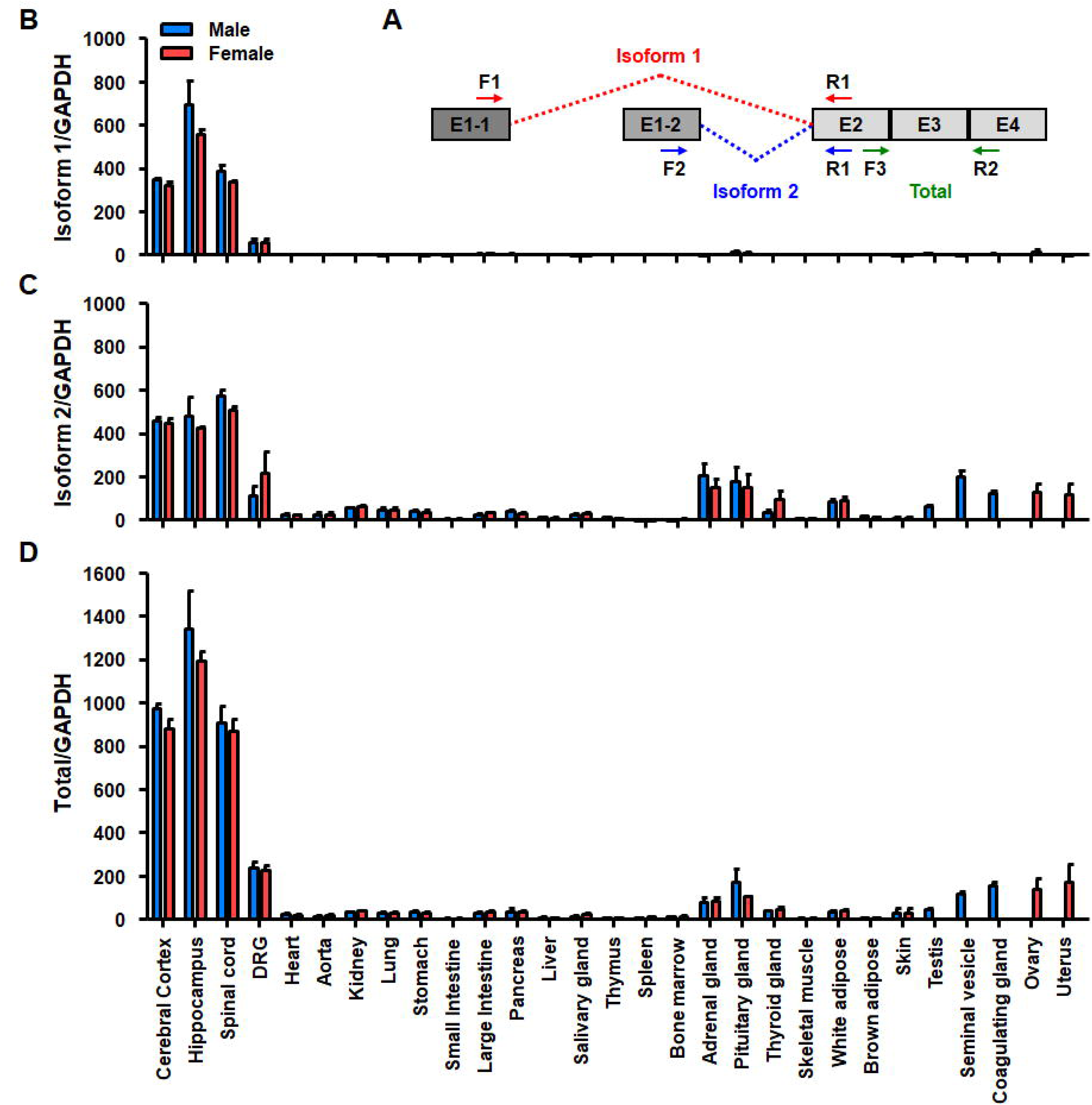
FAM19A5 transcript levels were measured by qRT□PCR. (A) A schematic diagram illustrating the PCR primer locations used to determine the FAM19A5 isoform 1 (Iso-1), isoform 2 (Iso-2), and total FAM19A5 transcripts. Iso-1 and iso-2 differ in the exon 1 sequence (E1-1 for iso-1 and E1-2 for iso-2) and share exons 2-4. Dashed lines indicate splicing junctions for iso-1 (red) and iso-2 (blue). Iso-1, iso-2, and total transcript levels were analyzed using F1-R1, F2-R2, and F3-R3 primers, respectively. (B-D) qRT□PCR analysis of FAM19A5 iso-1 (B), iso-2 (C) and total transcript (D) levels in the indicated tissues. The values are presented as the means ± standard errors of the means from three independent experiments.

**Table 1.**
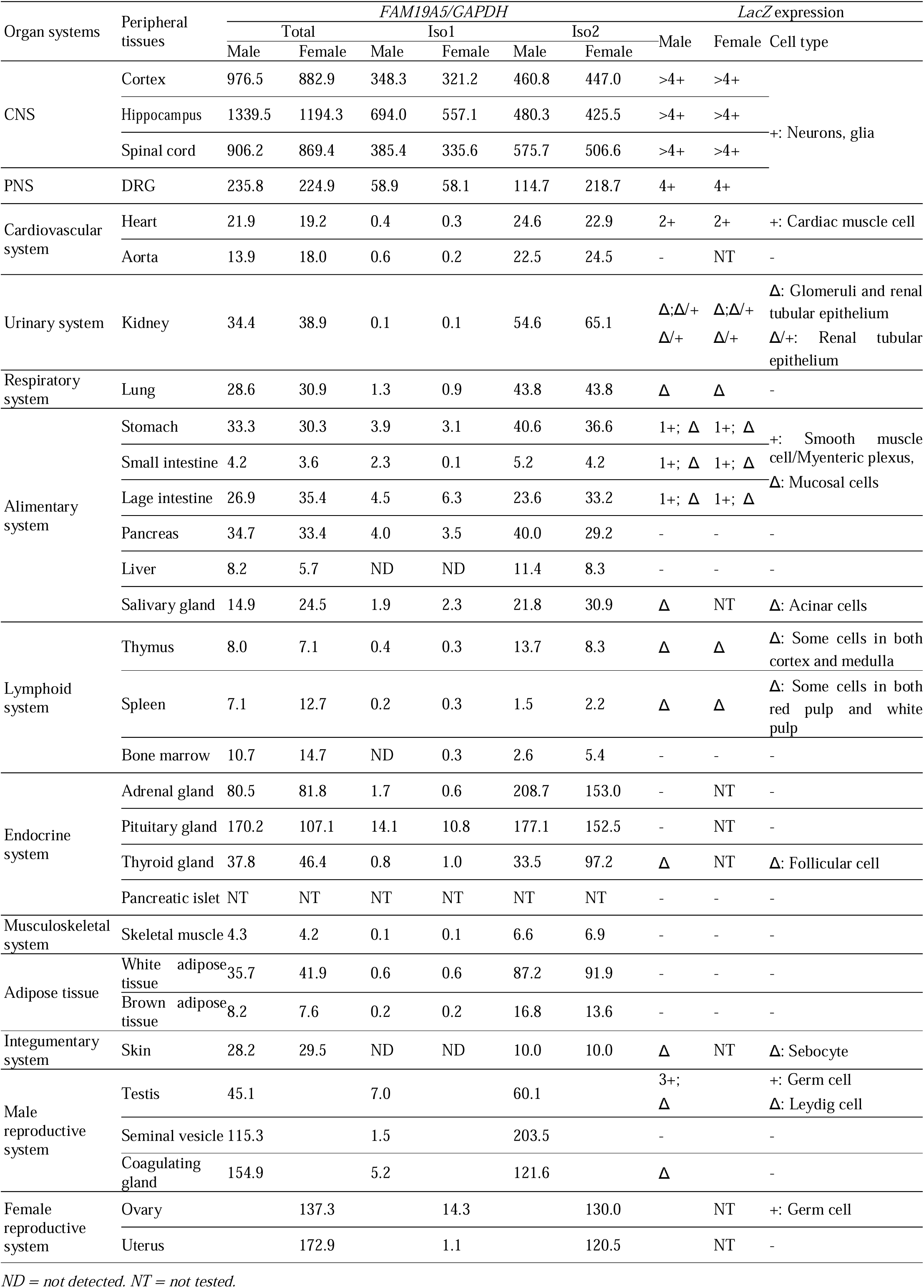
*FAM19A5* expression profile of mouse peripheral tissues using qRT□PCR and X□gal staining of wild-type and *FAM19A5-LacZ* knockin (+/+) mice.

Tissues from the central nervous system (CNS) exhibited the highest levels of isoform 1/2 and total FAM19A5 transcripts. The dorsal root ganglion (DRG), a representative peripheral nervous system (PNS) tissue, also exhibited high FAM19A5 transcript levels. For these CNS and PNS tissues, the FAM19A5 transcript levels were not significantly different between male and female mice (Fig. 1B, C, D, and Table 1).

In general, peripheral tissues exhibited very low levels of FAM19A5 transcripts, ranging between 1/100 and 1/10 of the total transcript levels found in CNS tissues. Notably, isoform 1 expression was negligibly low in all examined peripheral tissues. In contrast, isoform 2 was moderately expressed in reproductive tissues and endocrine tissues, including the thyroid, pituitary, and adrenal glands. Other tissues, including white and brown adipose tissue, exhibited low levels of total transcripts and isoform 2 (Fig. 1B, C, D, and Table 1).

### FAM19A5 expression in peripheral tissues from FAM19A5-LacZ KI mice

To identify the cell types that express the FAM19A5 transcript, we used X-gal staining of tissues from FAM19A5-LacZ KI mice (Shahapal et al., 2019). Positive X-gal staining in diverse cell types, including neurons, astrocytes, OPCs, and microglia, was consistent with the results of single-cell RNA sequencing of mouse and human brains (Zeisel et al., 2018; Karlsson et al., 2021; Sjöstedt et al., 2020; Mathy et al., 2019). Additionally, the X-gal signal intensity highly correlated with the FAM19A5 transcript levels measured by qRT□PCR (Shahapal et al., 2019).

In this study, we further identified peripheral tissues that express FAM19A5 using FAM19A5-LacZ KI mice. X-gal staining was performed on various peripheral tissues (Supplementary Table 1 and Supplementary Figs. 1–24). FAM19A5 promoter-driven X-gal signals were not observed in most tissues. The exceptions included subsets of cardiac muscle cells, smooth muscle cells or mesenteric plexuses of the gastrointestinal tract, inner cortical cells of the X-zone in the adrenal gland, myometrial smooth muscle cells in the uterus, and germ cells in the testis and ovary (Fig. 2 and Supplementary Figs. 1–24). The X-gal signal intensity in these cells was very faint to moderate compared to that in the hippocampus of the brain (Fig. 2). X-gal staining in white and brown adipose tissues was not detected, even though these tissues were reported to express high levels of FAM19A5 (Wang et al 2018).

**Fig. 2.**
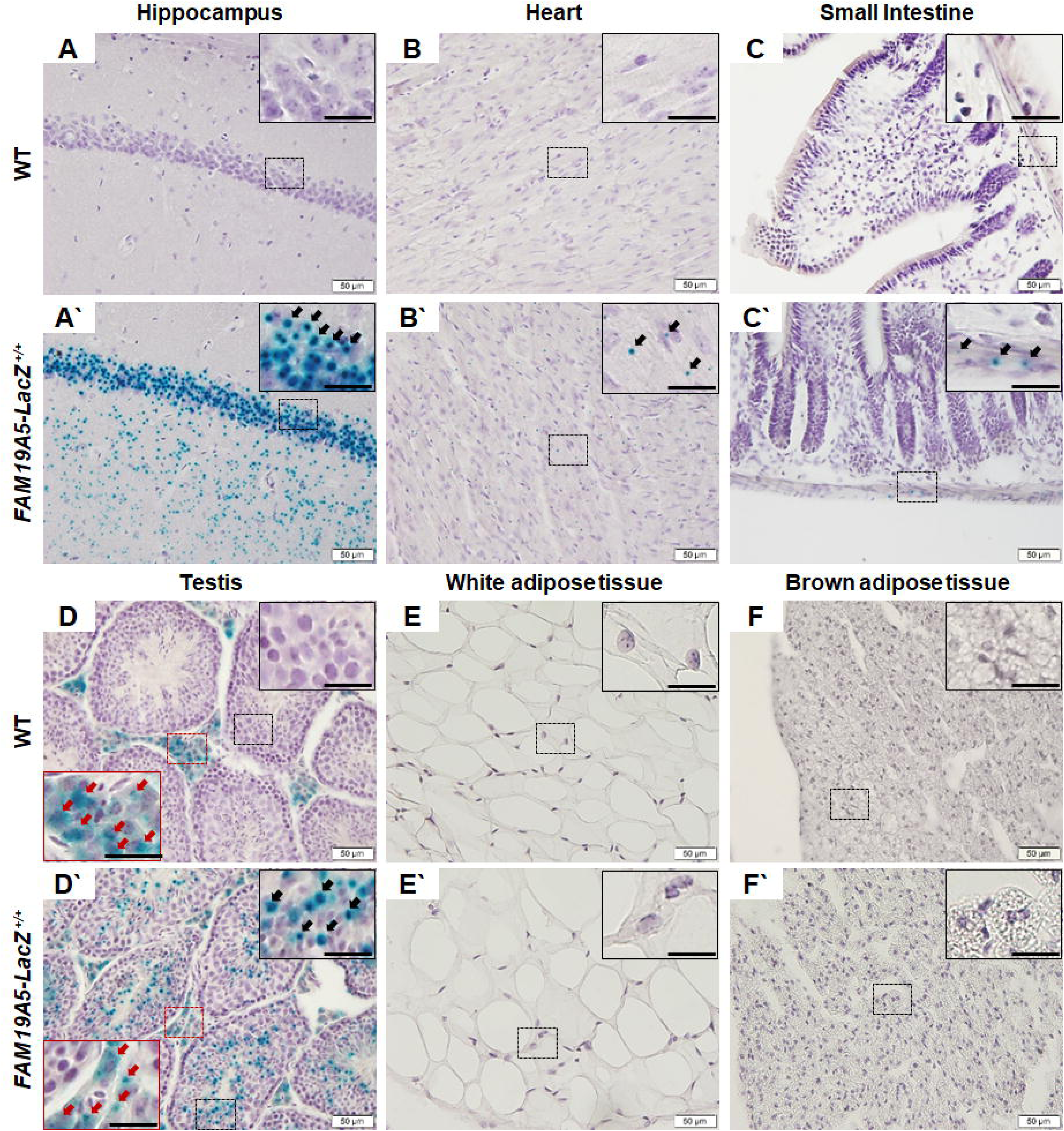
X-gal staining of peripheral tissues from wild-type and FAM19A5-LacZ knockin (+/+) mice. (A-F) Cryosections were stained with X-gal and counterstained with hematoxylin. Representative light photomicrographs of the brain hippocampus (A, A’), heart (B, B’), small intestine (C, C’), testis (D, D’), white adipose tissue (E, E’), and brown adipose tissue (F, F’) from wild-type (A-F) and FAM19A5-LacZ+/+ (A’-F’) mice. The images in the dashed boxes are magnified in the upper right. Dot-shaped blue precipitates were only present in the FAM19A5-LacZ+/+ samples, as indicated by the black arrows (A’, B’ C’ and D’). Dispersed precipitates were observed in both the WT and FAM19A5-LacZ+/+ samples and are indicated by red arrows (D and D’). The scale bars in the inset represent 20 μm.

The X-gal signals observed in WT mice were likely due to tissue-specific high endogenous β-gal expression (Johnson et al., 1986; Trifonov et al., 2016). The endogenous β-gal-driven X-gal-positive tissues included glomeruli and renal tubules in the kidney; individual cells in the gastrointestinal mucosa; alveolar macrophages; respiratory epithelia in the lung, thymus, and spleen immune cells; thyroid follicles; white and brown adipose tissues; sebaceous glands in the skin; and the coagulating gland epithelium, Leydig cells in the testis, endometrial glandular epithelium in the uterus, and granulosa and lutein cells in the ovary (Fig. 2, Supplementary Table 1, and Supplementary Figs. 1–24).

### FAM19A5 protein levels in peripheral tissues

FAM19A5 KO mice were generated with the CRISPR/Cas9 system (Fig. 3A, and B), and their FAM19A5 expression levels were measured (Fig. 3C, E, and F). qRT□PCR analysis indicated that the expression of the 1/2 isoform of FAM19A5 was not detectable in brain tissue from the FAM19A5 KO mice (Fig. 3C).

**Fig. 3.**
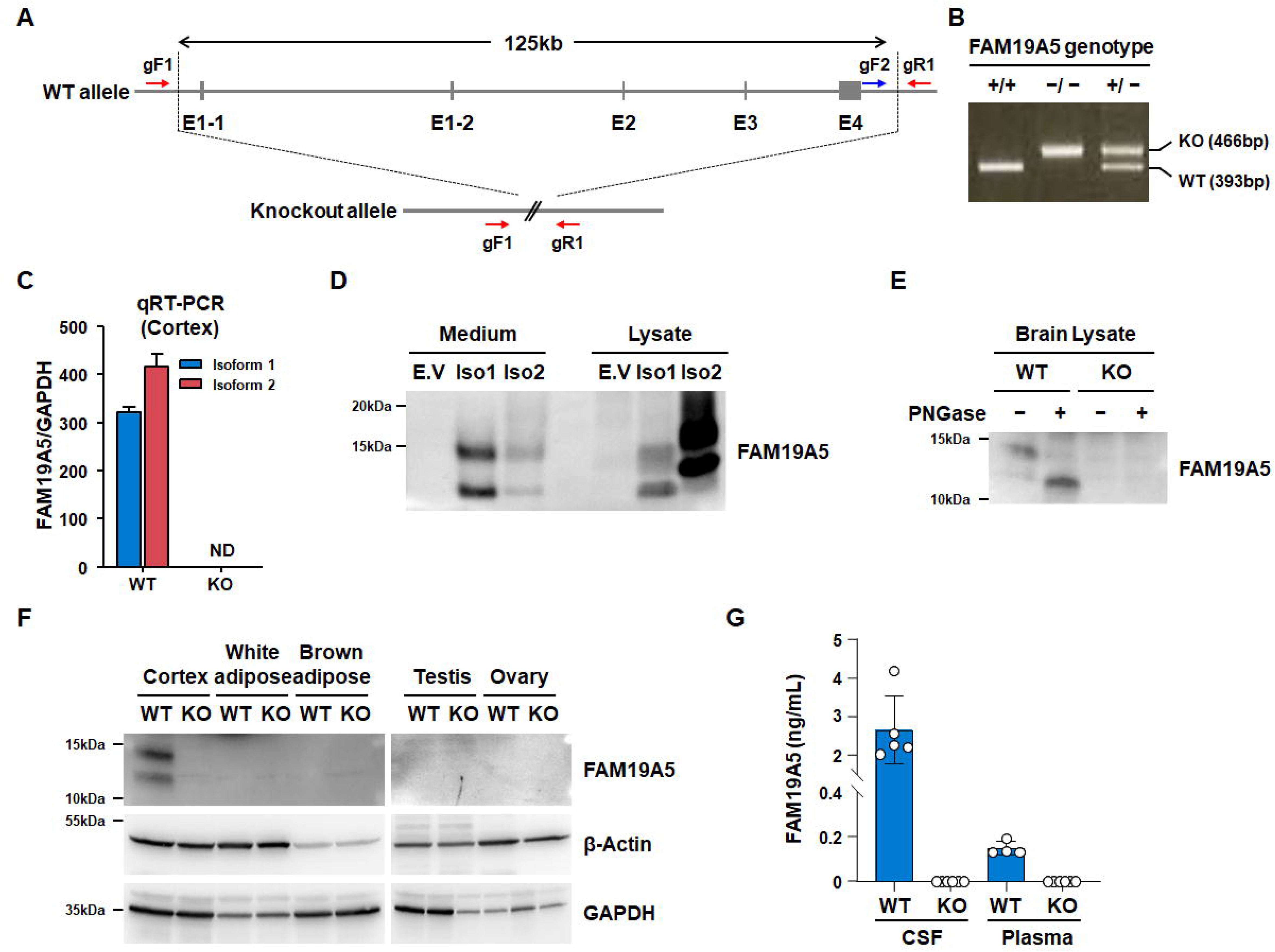
FAM19A5 protein levels in wild-type and FAM19A5 KO mice. (A) A schematic diagram illustrating the genome structures of the wild-type (WT) and FAM19A5 knockout (KO) alleles. The FAM19A5 gene from exon 1-1 (E1-1) to exon 4 (E4) was removed using the CRISPR/Cas9 system. The PCR primer locations used to determine the WT (gF1-gR1) and KO (gF2-gR1) alleles are indicated. (B) Genotyping of WT (+/+), heterozygous (+/-), and homozygous (-/-) FAM19A5 KO mice using genomic DNA. (C) qRT□PCR analysis of the FAM19A5 transcripts iso-1 and iso-2 in the cortex of WT and FAM19A5 KO mice. (D) Western blot analysis of FAM19A5 in culture medium and lysates from HEK293 cells expressing FAM19A5-Iso1 or FAM19A5-Iso2, respectively. EV: Empty vector. (E) Western blot analysis of FAM19A5 proteins extracted from the brains of WT and FAM19A5 KO mice. The protein samples were treated with PNGase to determine the presence of glycosylated forms of the FAM19A5 protein. (F) FAM19A5 protein levels in white and brown adipose tissues, testes, and ovaries of WT and FAM19A5 KO mice. β-actin and GAPDH were used as internal controls. (G) Quantitative analysis of FAM19A5. FAM19A5 levels in the CSF and plasma of WT and FAM19A5 KO mice.

The FAM19A5 proteins corresponding to the two FAM19A5 isoform transcripts differ in their N-terminal sequences. Isoform 1 produces a precursor protein that is cleaved at the end of the signal peptide to generate a mature, secreted, 89 amino acid protein. Isoform 2, however, can have two different fates. Isoform 2 may generate a protein that associates with the plasma membrane due to a hydrophobic α-helix at the N-terminus. Alternatively, the N-terminal sequence can also serve as a signal peptide, resulting in a secreted protein that is identical to the mature protein produced by isoform 1 (Wang et al 2018).

We investigated the FAM19A5 protein expression patterns in HEK293 cells expressing each type of FAM19A5 isoform transcript. The isoform 1 protein was largely detected in the culture media, whereas a small amount of isoform 2 protein was detected in the media (Fig. 3D). In contrast, isoform 2 protein was more prevalent in the cell lysate than isoform 1 protein. This observation indicates that the isoform 2 protein is less secreted than the isoform 1 protein. The molecular weights of the isoform 1 protein samples collected from the media and cell lysates were similar, indicating that they were identical. On the other hand, the molecular weight of isoform 2 protein from cell lysates is greater than that from media. This finding suggested that the N-terminal region of the isoform 2 protein from cell lysates was not cleaved. Instead, this N-terminal region can serve as a transmembrane domain (TMD) due to its long hydrophobic amino acid stretch, allowing the protein to associate with the plasma membrane (Fig. 3D).

We then investigated the presence of the FAM19A5 protein in mouse brain extracts. Western blotting showed the presence of both glycosylated and nonglycosylated forms of FAM19A5. After treating the brain extracts with PNGase, a deglycosylation enzyme, the intensity of the band corresponding to glycosylated FAM19A5 greatly decreased, while the intensity of the band corresponding to nonglycosylated FAM19A5 increased. Neither glycosylated nor nonglycosylated forms of the FAM19A5 protein were observed in the brain extracts from FAM19A5 KO mice (Fig. 3E).

Next, we measured the FAM19A5 protein levels in white and brown adipose tissues because the FAM19A5 expression patterns in these tissues are not well defined (Wang et al 2018). We also measured FAM19A5 levels in the testis and ovary because they exhibited moderate levels of the FAM19A5 transcript. None of the peripheral tissues exhibited detectable FAM19A5 protein levels, which correlated with the measured FAM19A5 transcript levels (Fig. 3F).

Notably, the FAM19A5 proteins observed in the mouse tissue samples are likely the secreted form of isoform 1. The isoform 2 protein, mainly observed in HEK293 cell lysates with a higher molecular weight than isoform 1, was not detected in either the brain or any peripheral tissues. Therefore, while the FAM19A5 isoform 2 transcript is expressed in the brain and peripheral tissues, protein production from this isoform is likely negligible under physiological conditions.

We investigated the plasma and CSF levels of FAM19A5 in wild-type and FAM19A5 KO mice using ELISA. In wild-type mice, the levels of FAM19A5 in CSF were detected at 2 to 4 ng/ml, whereas the levels in plasma were approximately 100 to 200 pg/ml, which is near the lowest level of the ELISA standard curve. In contrast, FAM19A5 was not detected in the CSF or plasma of FAM19A5 KO mice, validating the specificity of the ELISA method used in this study for FAM19A5 (Fig. 3G). This result further confirms the low levels of plasma FAM19A5 secreted from peripheral tissues.

### FAM19A5 does not interact with S1PR2

A previous study reported that FAM19A5 can regulate blood vessel smooth muscle function by interacting with S1PR2 (Wang et al., 2018). To investigate whether FAM19A5 elicits S1PR2-mediated G protein signaling, we used an SRE-luc assay system coupled to a HEK293G_qi_ stable cell line (Kim et al., 2014). Treatment with sphingosine-1-phosphate (S1P), a natural agonist of S1PR2, for 6 hours substantially increased SRE-luc activity, while treatment with FAM19A5 did not (Fig. 4A). These data indicate that the FAM19A5 protein cannot induce G protein-mediated signal transduction in S1PR2-expressing cells.

**Fig. 4.**
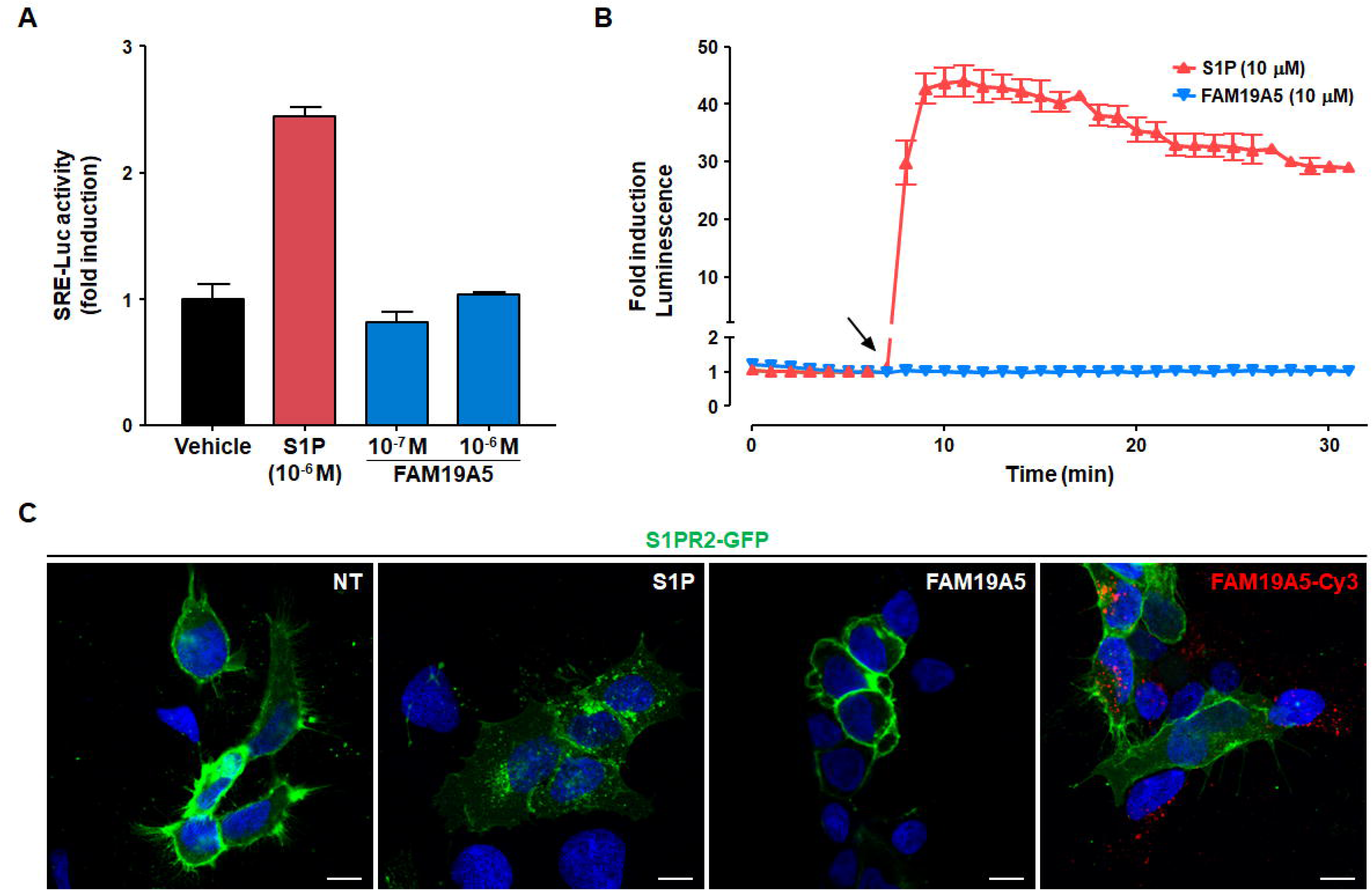
The FAM19A5 protein did not interact with S1PR2. (A) An SRE-luc reporter assay was used to determine S1PR2 receptor activation in response to S1P and FAM19A5. The SRE-luc reporter and S1PR2 genes were cotransfected into HEK293 cells that stably expressed the Gqi protein. Cells in serum-free medium were then treated with both S1P and FAM19A5 for 6 h and subjected to a luciferase assay. (B) Nanoluc reporter assay for β-arrestin recruitment to S1PR2. The S1PR2-LgBit and β-arrestin-SmBit constructs were cotransfected into HEK293 cells. In a live-cell system, luminescence was determined before and after treatment with S1P and FAM19A5. The data are presented as the means ± standard errors of the means from three independent experiments. (C) Internalization of S1PR2 in response to S1P and FAM19A5. HEK293 cells were transfected with the S1PR2-GFP construct. The cells were cultured under serum-free conditions for 16 hr and then treated with S1P, naïve FAM19A5, and FAM19A5-Cy3 for 30 min. The cellular locations of S1PR2-GFP in the presence of the ligands were determined using a confocal microscope. The scale bar represents 10 μm.

Since some biased agonists induce β-arrestin recruitment to S1PR2 independently of G-protein-mediated signaling (Kim et al., 2014), we explored whether FAM19A5 increased the interaction between S1PR2 and β-arrestin using the NanoLuc luciferase assay (Reyes-Alcaraz et al., 2018; Reyes-Alcaraz et al., 2019). S1P induced an immediate increase in luciferase activity, indicating an interaction between S1PR2 and β-arrestin. Conversely, FAM19A5 had no effect (Fig. 4B).

Next, we investigated the effect of FAM19A5 on the internalization of S1PR2. S1PR2 is primarily localized to the plasma membrane in the absence of ligand treatment. After treatment with S1P, the cellular internalization of S1PR2-GFP increased. However, treatment with FAM19A5 did not result in S1PR2-GFP internalization. Furthermore, FAM19A5-Cy3 was internalized regardless of S1PR2 expression (Fig. 4C). Collectively, these data indicate that FAM19A5 cannot induce either β-arrestin recruitment to S1PR2 or ligand-stimulated internalization of S1PR2.

## DISCUSSION

The quantitative levels of total FAM19A5 transcripts, isoform 1, and isoform 2 were investigated in a wide variety of peripheral tissues and compared to the levels in specific brain regions. In general, FAM19A5 transcript levels in peripheral tissues were very low compared to those in brain tissues. We hypothesize that isoform 2 is the major form of FAM19A5 in peripheral tissues, with isoform 1 being minimally expressed. The isoform 1 transcript produces a mature secretory protein, while the isoform 2 transcript may primarily produce an uncleaved plasma membrane-bound protein. However, immunoblotting of brain and peripheral tissues revealed the absence of isoform 2 transcript-derived uncleaved FAM19A5 protein. Therefore, the soluble secretory form of FAM19A5 is produced in much lower quantities in peripheral tissues than in the brain.

Similar to the low levels of FAM19A5 transcripts measured in peripheral tissues, FAM19A5 promoter-driven X-gal signals were also very faint or negligible in peripheral tissues compared to the signal in the brain. Only a few samples exhibited faint to moderate X-gal signals, including cardiac muscle cells, gastrointestinal cells in the muscularis externa, and germ cells in the testis and ovary. We did not detect X-gal signals in any other tissues, including white adipose tissue and the thyroid and adrenal glands. Because both the X-gal intensity and the total transcript levels were very low in peripheral tissues compared to those in the brain, we concluded that the levels of the FAM19A5 protein produced in these cells/tissues are likely very low. The functional relevance of these low-level proteins in peripheral tissues needs further investigation.

It is intriguing that no FAM19A5 promoter-driven X-gal signals were observed in adipose tissue cells, even though these tissues are reported to produce significantly high levels of both FAM19A5 transcripts and protein (Wang et al., 2018). Furthermore, qRT□PCR results demonstrated that transcript levels measured in white and brown adipose tissues were significantly lower than those measured in brain tissues. Our data are in agreement with the Human Protein Atlas Database, which includes three independent datasets (HPA, GTEx, and FANTOM5) showing that FAM19A5 transcript levels in human adipose tissue are approximately 1/100 to 1/50 of those found in the human brain (Uhlén et al., 2015; GTEx Consortium, 2015; Lizio et al., 2015). These datasets also show extremely low levels of FAM19A5 transcripts in peripheral tissues and low-to-moderate expression levels in female and male reproductive tissues. FAM19A5 expression in WT and FAM19A5 KO mice was also consistent with the available data. FAM19A5 protein in adipose and reproductive tissues was below detectable levels, but its expression in brain tissue was high.

We were unable to observe X-gal signals in the bone marrow, thymus, or spleen, which are blood cell-producing organs. Thus, the low levels of FAM19A5 transcripts in peripheral tissues raise questions about the origin of FAM19A5 in the blood. Recently, blood FAM19A5 levels were measured in human (Lee et al., 2019) and mouse samples (Wang et al., 2018). The blood levels of FAM19A5 measured by ELISA ranged from 100 pg/ml for human samples (Lee et al., 2019) to 400 ng/ml for mouse samples (Wang et al., 2018). Our recently validated ELISA, however, showed that plasma concentrations of the FAM19A5 protein are usually less than 500 pg/ml in both humans and mice, while its concentrations in CSF are much greater, ranging from 2 to 10 ng/ml (Kim et al., 2023).

In conclusion, our research suggested that plasma FAM19A5 levels may not be sufficient to exert a functional effect on peripheral tissues under physiological conditions. Moreover, FAM19A5 does not appear to directly interact with the S1PR2 receptor to trigger G protein activation or β-arrestin recruitment, leading to receptor internalization. Further research is needed to understand the functional relevance of the low levels of the FAM19A5 protein in peripheral tissues under both physiological and pathological conditions.

## ACKNOWLEDGMENTS

This research was supported by grants from Neuracle Science Co., Ltd and the Research Program of the National Research Foundation of Korea (NRF□2020M3E5D9080794), which is funded by the Korea Government (MSIT).

## AUTHOR CONTRIBUTIONS

JY.S., HY.K., EH.C., and WS.L. wrote the manuscript. HY.K., EH.C., EB.C., YN.L., MH.L., N.H., and ST.O. performed the experiments. JY.S., HY.K., and EH.C. analyzed the data. A.S., HJ.Y., A.RA., YW.J., YR.L., and WK.K. provided reagents and samples. SY.S., JK.L., and JI.H provided expertise and feedback. JY.S. supervised the research and secured funding.

## CONFLICTS OF INTEREST

HY.K., EH.C., EB.C., YW.J., YR.L., MH.L., N.H., ST.O., JK.L., WS.L., WK.K., and JY.S. are employed by Neuracle Science Co., Ltd. A.S., HJ.Y., A.RA., YW.J., MH.L., WK.K., and JY.S. are shareholders of Neuracle Science Co., Ltd. The other authors have no conflicts of interest to declare.

## Notes

### Summary of Updates

Major changes to the manuscript include the following: 1. We have added western blotting experiments to investigate the expression patterns of FAM19A5 isoform 1 and isoform 2 in culture medium and cell lysate. The results showed that isoform 1 protein was detected in the culture medium, whereas isoform 2 protein was more prevalent in the cell lysate. Furthermore, we confirmed that the isoform 2 protein observed in the cell lysate exists without cleavage at the N-terminal region. 2. We conducted FAM19A5 ELISA in the CSF and plasma of wild-type and FAM19A5 knock-out mice. This experiment confirmed the low levels of plasma FAM19A5 secreted from peripheral tissues. 3. The content related to Figure 5 from the previous version of the manuscript has been removed as it was deemed inappropriate for the revised manuscript.

## REFERENCES

Conklin, B.R., Farfel, Z., Lustig, K.D., Julius, D., and Bourne, H.R. (1993). Substitution of three amino acids switches receptor specificity of Gq alpha to that of Gi alpha. Nat. 363, 274–276.

GTEx Consortium (2015). Human genomics. The Genotype-Tissue Expression (GTEx) pilot analysis: multitissue gene regulation in humans. Science 348, 648–660.

Huang, S., Zheng, C., Xie, G., Song, Z., Wang, P., Bai, Y., Chen, D., Zhang, Y., Lv, P., Liang, W., et al. (2021). FAM19A5/TAFA5, a novel neurokine, plays a crucial role in depressive-like and spatial memory-related behaviors in mice. Mol. Psychiatry 26, 2363–2379.

Jeong, I., Yun, S., Shahapal, A., Cho, E.B., Hwang, S.W., Seong, J.Y., and Park, H.C. (2021). FAM19A5l affects mustard oil-induced peripheral nociception in zebrafish. Mol. Neurobiol. 58, 4770–4785.

Johnson, W.G., Hong, J.L., and Knights, S.M. (1986). Variation in ten lysosomal hydrolase enzyme activities in inbred mouse strains. Biochem. Genet. 24, 891–909.

Karlsson, M., Zhang, C., Méar, L., Zhong, W., Digre, A., Katona, B., Sjöstedt, E., Butler, L., Odeberg, J., Dusart, P., et al. (2021). A single-cell type transcriptomics map of human tissues. Sci. Adv. 7, e2169.

Kim, D.K., Yun, S., Son, G.H., Hwang, J.I., Park, C.R., Kim, J.I., Kim, K., Vaudry, H., and Seong, J.Y. (2014). Coevolution of the spexin/galanin/kisspeptin family: Spexin activates galanin receptor type II and III. Endocrinology 155, 1864–1873.

Kim, H.B., Yoo, S., Kwak, H., Ma, S.X., Kim, R., Lee, M., Ha, N., Pyo, S., Kwon, S.G., Cho, E.H., et al. (2023). Inhibition of FAM19A5 restores synaptic loss and improves cognitive function in mouse models of Alzheimer’s disease. bioRxiv, 2023.11.22.568357.

Lee, J.H., Park, J.H., Nam, T.W., Seo, S.M., Kim, J.Y., Lee, H.K., Han, J.H., Park, S.Y., Choi, Y.K., and Lee, H.W. (2018). Differences between immunodeficient mice generated by classical gene targeting and CRISPR/Cas9-mediated gene knockout. Transgenic Res. 27, 241–251.

Lee, Y.B., Hwang, H.J., Kim, J.A., Hwang, S.Y., Roh, E., Hong, S.H., Choi, K.M., Baik, S.H., and Yoo, H. J. (2019). Association of serum FAM19A5 with metabolic and vascular risk factors in human subjects with or without type 2 diabetes. Diab. Vasc. Dis. Res., 16, 530–538.

Lizio, M., Harshbarger, J., Shimoji, H., Severin, J., Kasukawa, T., Sahin, S., Abugessaisa, I., Fukuda, S., Hori, F., Ishikawa-Kato, S., et al. (2015). Gateways to the FANTOM5 promoter level mammalian expression atlas. Genome Biol. 16, 22.

Mathys, H., Davila-Velderrain, J., Peng, Z., Gao, F., Mohammadi, S., Young, J.Z., Menon, M., He, L., Abdurrob, F., Jiang, X., et al. (2019). Single-cell transcriptomic analysis of Alzheimer’s disease. Nat. 570, 332–337.

Mountford, P., Zevnik, B., Düwel, A., Nichols, J., Li, M., Dani, C., Robertson, M., Chambers, I., and Smith, A. (1994). Dicistronic targeting constructs: reporters and modifiers of mammalian gene expression. Proc. Natl. Acad. Sci. USA 91, 4303–4307.

Reyes-Alcaraz, A., Lee, Y.N., Yun, S., Hwang, J.I., and Seong, J.Y. (2018). Conformational signatures in β-arrestin2 reveal natural biased agonism at a G-protein-coupled receptor. Commun. Biol. 1, 128.

Reyes-Alcaraz, A., Lee, Y.N., Yun, S., Hwang, J.I., and Seong, J.Y. (2019). Monitoring GPCR-β-arrestin1/2 interactions in real time living systems to accelerate drug discovery. J. Vis. Exp. 148, e59994.

Shahapal, A., Cho, E.B., Yong, H.J., Jeong, I., Kwak, H., Lee, J.K., Kim, W., Kim, B., Park, H. C., Lee, W.S., et al. (2019). FAM19A5 expression during embryogenesis and in the adult traumatic brain of FAM19A5-LacZ knock-in mice. Front. Neurosci. 13, 917.

Shahapal, A., Park, S., Yoo, S., and Seong., J.Y. (2024). Partial FAM19A5 deficiency in mice leads to disrupted spine maturation, hyperactivity and an altered fear response. bioRxiv. 2024.04.29.591582.

Sjöstedt, E., Zhong, W., Fagerberg, L., Karlsson, M., Mitsios, N., Adori, C., Oksvold, P., Edfors, F., Limiszewska, A., Hikmet, F., et al. (2020). An atlas of the protein-coding genes in the human, pig, and mouse brain. Science 367, e5947.

Tom Tang, Y., Emtage, P., Funk, W.D., Hu, T., Arterburn, M., Park, E.E., and Rupp, F. (2004). TAFA: a novel secreted family with conserved cysteine residues and restricted expression in the brain. Genomics 83, 727–734.

Trifonov, S., Yamashita, Y., Kase, M., Maruyama, M., and Sugimoto, T. (2016). Overview and assessment of the histochemical methods and reagents for the detection of β-galactosidase activity in transgenic animals. Anat. Sci. Int. 91, 56–67.

Uhlén, M., Fagerberg, L., Hallström, B.M., Lindskog, C., Oksvold, P., Mardinoglu, A., Sivertsson, Å., Kampf, C., Sjöstedt, E., Asplund, A., et al. (2015). Proteomics. Tissue-based map of the human proteome. Science 347, 1260419.

Wang, Y., Chen, D., Zhang, Y., Wang, P., Zheng, C., Zhang, S., Yu, B., Zhang, L., Zhao, G., Ma, B., et al. (2018). Novel adipokine, FAM19A5, inhibits neointima formation after injury through sphingosine-1-phosphate receptor 2. Circ. 138, 48–63.

Woo, J., Kwon, S.K., Choi, S., Kim, S., Lee, J.R., Dunah, A.W., Sheng, M., and Kim, E. (2009). Trans-synaptic adhesion between NGL-3 and LAR regulates the formation of excitatory synapses. Nat. Neurosci. 12, 428–437.

