## Supplementary materials for "Is FAM19A5 an adipokine? Peripheral FAM19A5 in wild-type, FAM19A5 knockout, and LacZ knockin mice"

**Is FAM19A5 an adipokine? Peripheral FAM19A5 in wild-type, FAM19A5 knock-out, and LacZ knock-in mice**

**Supplementary Table 1. X-gal staining for peripheral tissues**

| Peripheral organ/tissue | | | Group | | | | | | | |
| --- | --- | --- | --- | --- | --- | --- | --- | --- | --- | --- |
|  |  |  | WT | | | *Hetero*  (+/-) | *Homozygote*  (+/+) | | | |
|  |  |  | Male#1 | Male#2 | Female#1 | Male#1 | Male#1 | Male#2 | Male#3 | Female #1 |
| Heart | Cardiac muscle cell | | - | - | N.T. | 1+ | 2+ | 1+ | 1+ | 2+ |
| Stomach | Mucosa | Gastric gland | Δ | Δ | N.T. | Δ | Δ | Δ | Δ | N.T. |
|  | Submucosa | Connective tissue | - | - | N.T. | - | - | - | - | N.T. |
|  | Muscularis  externa | Smooth muscle & Mesenteric plexus | - | - | N.T. | 1+ | 1+ | - | 1+ | 1+ |
| Small intestine | Mucosa | Villus & Crypt | - | Δ | N.T. | - | - | - | - | - |
|  | Submucosa | Connective tissue | - | - | N.T. | - | - | - | - | - |
|  | Muscularis externa | Smooth muscle & mesenteric plexus | - | - | N.T. | 1+ | 1+ | 1+ | 1+ | 1+ |
| Large intestine | Mucosa | Villus & Crypt | Δ | Δ | N.T. | Δ | Δ | Δ | Δ | Δ |
|  | Submucosa | Connective tissue | - | - | N.T. | - | - | - | N.T. | - |
|  | Muscularis externa | Smooth muscle & Mesenteric plexus | - | - | N.T. | 1+ | 1+ | 1+ | N.T. | 1+ |
| Testis | Seminiferous tubule | Sertoli cell &  Germ cell | - | - |  | 2+ | N.T. | 3+ | 3+ |  |
|  | Interstitium | Leydig cell | Δ | Δ |  | Δ | N.T. | Δ | Δ |  |
| Ovary | Ovarian follicle | Oocyte |  |  | - |  |  |  |  | 1+ |
|  |  | Granulosa cell |  |  | Δ |  |  |  |  | Δ |
|  | Corpus luteun | Lutein cell |  |  | Δ |  |  |  |  | Δ |
| Uterus | Endometrium | Endometrial gland |  |  | Δ |  |  |  |  | Δ |
|  |  | Endometrial epithelium |  |  | Δ |  |  |  |  | Δ |
|  |  | Endometrial stroma |  |  | - |  |  |  |  | - |
|  | Myometrium | Smooth muscle cell |  |  | - |  |  |  |  | 1+ |
| Adrenal gland | Cortex | | - | N.T. | - | - | - | N.T. | N.T. | - |
|  | Medulla | | - | N.T. | - | - | - | N.T. | N.T. | - |
|  | X-Zone | | N.T. | N.T. | - | N.T. | N.T. | N.T. | N.T. | 1+ |
| Aorta | Endothelium | | N.T. | - | N.T. | N.T. | N.T. | - | - | N.T. |
|  | Smooth muscle & Connective tissue | | N.T. | - | N.T. | N.T. | N.T. | - | - | N.T. |
| Kidney | Glomerulus | | Δ | Δ | N.T. | Δ | Δ | Δ | Δ | Δ |
|  | Renal tubule | | Δ | Δ | N.T. | Δ | Δ/+ | Δ | Δ | Δ/+ |
| Lung | Pseudostratified columnar epithelium | | Δ | N.T. | N.T. | Δ | Δ | N.T. | N.T. | - |
|  | Pneumocyte | | - | N.T. | N.T. | - | - | N.T. | N.T. | - |
|  | Alveolar macrophage | | Δ | N.T. | N.T. | Δ | Δ | N.T. | N.T. | - |
| Pancreas | Exocrine gland | | - | N.T. | N.T. | - | - | N.T. | N.T. | - |
|  | Endocrine gland (pancreatic islet) | | - | N.T. | N.T. | - | - | N.T. | N.T. | - |
| Liver | Hepatocyte | | - | N.T. | N.T. | - | - | N.T. | N.T. | - |
|  | Bile duct | | - | N.T. | N.T. | - | - | N.T. | N.T. | - |
|  | Central vein | | - | N.T. | N.T. | - | - | N.T. | N.T. | - |
|  | Portal vein | | - | N.T. | N.T. | - | - | N.T. | N.T. | - |
| Submandibular  gland | Serous acinus | | N.T. | Δ | N.T. | N.T. | N.T. | Δ | Δ | N.T. |
|  | Mucous acinus | | N.T. | Δ | N.T. | N.T. | N.T. | Δ | Δ | N.T. |
|  | Connective tissue | | N.T. | - | N.T. | N.T. | N.T. | - | - | N.T. |
| Sublingual  gland | Mucous acinus | | N.T. | Δ | N.T. | N.T. | N.T. | N.T. | Δ | N.T. |
|  | Connective tissue | | N.T. | - | N.T. | N.T. | N.T. | N.T. | - | N.T. |
| Parotid gland | Serous acinus | | N.T. | Δ | N.T. | N.T. | N.T. | Δ | Δ | N.T. |
|  | Connective tissue | | N.T. | - | N.T. | N.T. | N.T. | - | - | N.T. |
| Thymus | Cortex | | Δ | N.T. | N.T. | Δ | Δ | N.T. | N.T. | Δ |
|  | Medulla | | Δ | N.T. | N.T. | Δ | Δ | N.T. | N.T. | Δ |
| Spleen | White pulp | | Δ | N.T. | N.T. | Δ | Δ | N.T. | N.T. | Δ |
|  | Red pulp | | Δ | N.T. | N.T. | Δ | Δ | N.T. | N.T. | Δ |
| Bone marrow | Hematopoietic cell | | - | N.T. | N.T. | - | - | N.T. | N.T. | - |
| Pituitary gland | Pars nervosa | | N.T. | - | N.T. | N.T. | N.T. | - | - | N.T. |
|  | Pars intermediate | | N.T. | - | N.T. | N.T. | N.T. | - | - | N.T. |
|  | Pars distalis | | N.T. | - | N.T. | N.T. | N.T. | - | N.T. | N.T. |
| Thyroid gland | Follicular cell & Parafollicular cell | | N.T. | Δ | N.T. | N.T. | N.T. | N.T. | Δ | N.T. |
|  | Connective tissue | | N.T. | - | N.T. | N.T. | N.T. | N.T. | - | N.T. |
| Skeletal muscle | | | - | N.T. | N.T. | - | - | N.T. | N.T. | - |
| White adipose  tissue | White adipocyte | | Δ | N.T. | N.T. | N.T. | Δ | N.T. | N.T. | - |
|  | Connective tissue | | - | N.T. | N.T. | N.T. | - | N.T. | N.T. | - |
| Brown adipose  tissue | Brown adipocyte | | Δ | N.T. | N.T. | - | - | N.T. | N.T. | - |
|  | Connective tissue | | - | N.T. | N.T. | - | - | N.T. | N.T. | - |
| Skin | Epidermis | Stratified squamous epithelium | N.T. | - | N.T. | N.T. | N.T. | - | - | N.T. |
|  | Dermis | Hair follicle | N.T. | - | N.T. | N.T. | N.T. | - | - | N.T. |
|  |  | Sebaceous gland | N.T. | Δ | N.T. | N.T. | N.T. | Δ | Δ | N.T. |
|  |  | Connective tissue | N.T. | - | N.T. | N.T. | N.T. | - | - | N.T. |
|  | Hypodermis | White adipose tissue | N.T. | - | N.T. | N.T. | N.T. | - | - | N.T. |
|  |  | Striated muscle cell | N.T. | - | N.T. | N.T. | N.T. | - | - | N.T. |
|  |  | Peripheral nerve | N.T. | - | N.T. | N.T. | N.T. | - | - | N.T. |
| Seminal vesicle | Pseudostratified columnar epithelium | | - | - | N.T. | N.T. | - | - | - | N.T. |
| Coagulating  gland | Pseudostratified columnar epithelium | | Δ | Δ | N.T. | N.T. | N.T. | Δ | Δ | N.T. |

Ten-week-old adult wild-type littermates (2 males and 1 female) and *FAM19A5-LacZ* KI heterozygote (1 male) and homozygote (3 males and 1 female) mice were used for X-gal staining. Criteria for grading X-gal signal were described in the Materials and Methods section. N.T.: Not Tested.

**Supplementary Fig. 1. Heart X-gal signal.**

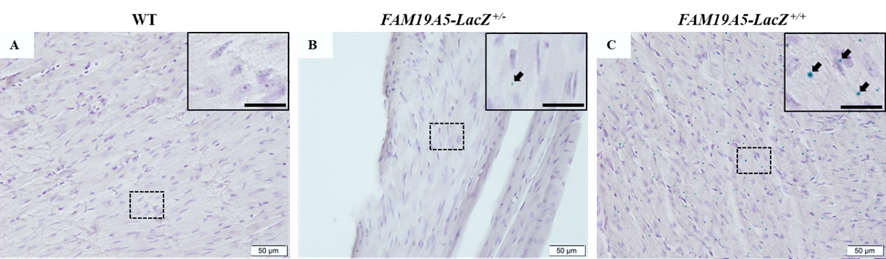

Representative light photomicrographs for heart cryosection of *(A)* wild type (WT, male #1), *(B) FAM19A5-LacZ*^+/-^ (Heterozygote, male #1), and *(C)* *FAM19A5-LacZ*^+/+^ (Homozygote, male #1). Cryosections were stained with X-gal solution and counter-stained with hematoxylin. Image in the dashed box is magnified in the inset. Black arrows indicate punctate blue precipitates in cardiac muscle cells observed only in *FAM19A5-LacZ* KI mice but not in WT mice. Scale bars in the inset represent 20 μm.

**Supplementary Fig. 2. Stomach X-gal signal.**

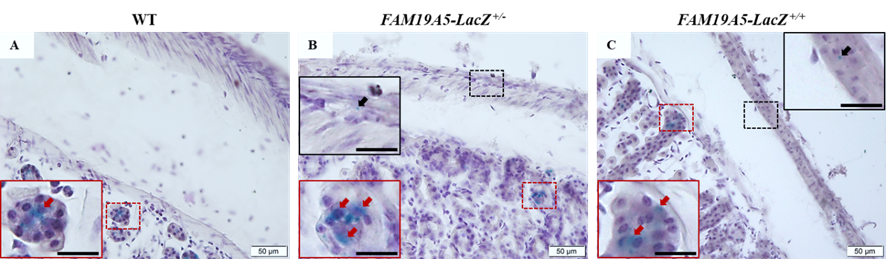

Representative light photomicrographs of stomach cryosection of *(A)* wild type (WT, male #1), *(B) FAM19A5-LacZ*^+/-^ (Heterozygote, male #1), and *(C)* *FAM19A5-LacZ*^+/+^ (Homozygote, male #3). Cryosections were stained with X-gal solution and counter-stained with hematoxylin. Image in the dashed box is magnified in the inset with the same color as the dashed box. Black arrows indicate punctate blue precipitates in smooth muscle cells or myenteric plexuses observed only in *FAM19A5-LacZ* KI mice but not in WT mice. Red arrows indicate dispersed blue precipitates in gastric glands observed in both WT and *FAM19A5-LacZ* KI mice. Scale bars in the inset represent 20 μm.

**Supplementary Fig. 3. Small intestine X-gal signal.**

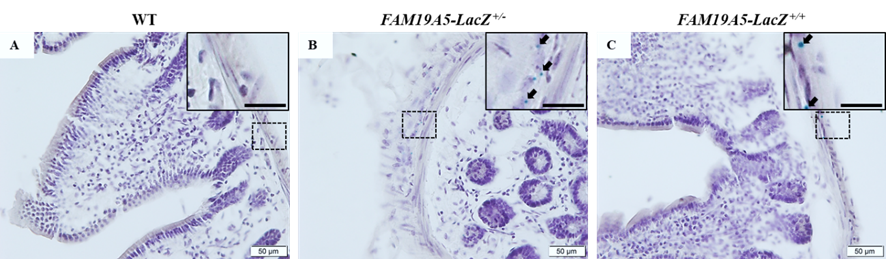

Representative light photomicrographs of small intestine cryosection of *(A)* wild type (WT, male #1), *(B) FAM19A5-LacZ*^+/-^ (Heterozygote, male #1), and *(C)* *FAM19A5-LacZ*^+/+^ (Homozygote, male #1). Cryosections were stained with X-gal solution and counter-stained with hematoxylin. Image in the dashed box is magnified in the inset with the same color as the dashed box. Black arrows indicate punctate blue precipitates in smooth muscle cells or myenteric plexuses observed only in *FAM19A5-LacZ* KI mice but not in WT mice. Scale bars in the inset represent 20 μm.

**Supplementary Fig. 4. Large intestine X-gal signal.**

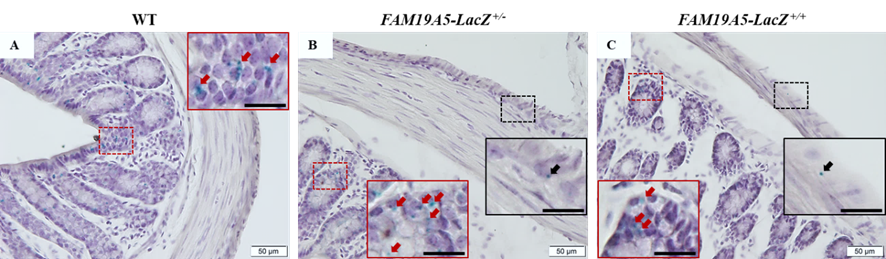

Representative light photomicrographs of large intestine cryosection of *(A)* wild type (WT, male #1), *(B) FAM19A5-LacZ*^+/-^ (Heterozygote, male #1), and *(C)* *FAM19A5-LacZ*^+/+^ (Homozygote, male #1). Cryosections were stained with X-gal solution and counter-stained with hematoxylin. Image in the dashed box is magnified in the inset with the same color as the dashed box. Black arrows indicate punctate blue precipitates in smooth muscle cells or myenteric plexuses observed only in *FAM19A5-LacZ* KI mice but not in WT mice. Red arrows indicate punctate blue precipitates in mucosa observed in both WT and *FAM19A5-LacZ* KI mice. Scale bars in the inset represent 20 μm.

**Supplementary Fig. 5. Testis X-gal signal.**

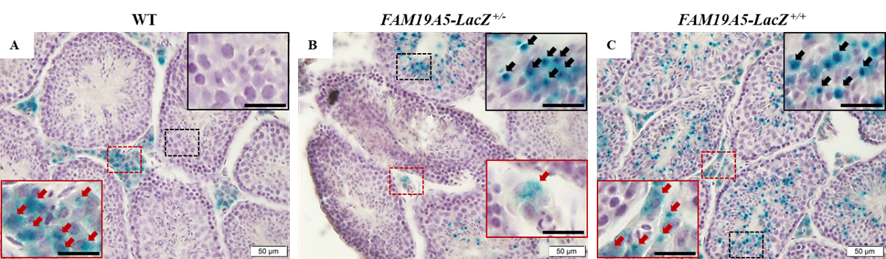

Representative light photomicrographs of testis cryosection of *(A)* wild type (WT, male #2), *(B) FAM19A5-LacZ*^+/-^ (Heterozygote, male #1), and *(C)* *FAM19A5-LacZ*^+/+^ (Homozygote, male #2). Cryosections were stained with X-gal solution and counter-stained with hematoxylin. Image in the dashed box is magnified in the inset with the same color as the dashed box. Black arrows indicate punctate blue precipitates in germ cells observed only in *FAM19A5-LacZ* KI mice but not in WT mice. Red arrows indicate dispersed blue precipitates in interstitial cells (Leydig cells) observed in both WT and *FAM19A5-LacZ* KI mice. Scale bars in the inset represent 20 μm.

**Supplementary Fig. 6. Ovary X-gal signal.**

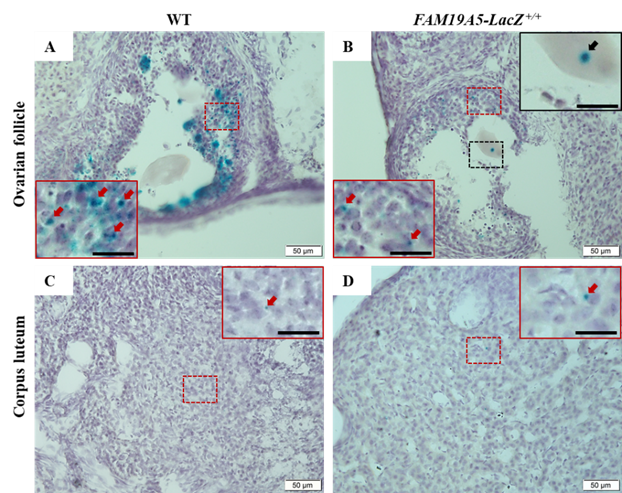

Representative light photomicrographs of ovary cryosection of *(A, C)* wild type (WT, female #1) and *(B, D) FAM19A5-LacZ*^+/+^ (Homozygote, female #1). Ovarian follicle (A-B) and corpus luteum (C-D) were analyzed. Cryosections were stained with X-gal solution and counter-stained with hematoxylin. Image in the dashed box is magnified in the inset with the same color as the dashed box. Black arrows indicate punctate blue precipitates in oocytes observed in *FAM19A5-LacZ* KI mice but not in WT mice. Red arrows indicate punctate/dispersed blue precipitates in granulosa and lutein cells observed in both WT and *FAM19A5-LacZ*^+/+^ mice. Scale bars in the inset represent 20 μm.

**Supplementary Fig. 7. Uterus X-gal signal.**

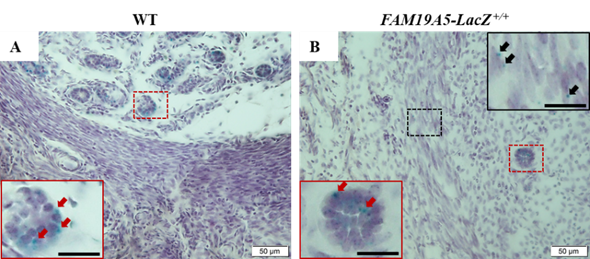

Representative light photomicrographs of uterus cryosection of *(A)* wild type (WT, female #1) and *(B) FAM19A5-LacZ*^+/+^ (Homozygote, female #1). Cryosections were stained with X-gal solution and counter-stained with hematoxylin. Image in the dashed box is magnified in the inset with the same color as the dashed box. Black arrows indicate punctate blue precipitates in myometrial smooth muscle cells observed only in *FAM19A5-LacZ* KI mice but not in WT mice. Red arrows indicate punctate/dispersed blue precipitates in endometrial gland observed in both WT and *FAM19A5-LacZ*^+/+^ mice. Scale bars in the inset represent 20 μm.

**Supplementary Fig. 8. Adrenal gland X-gal signal.**

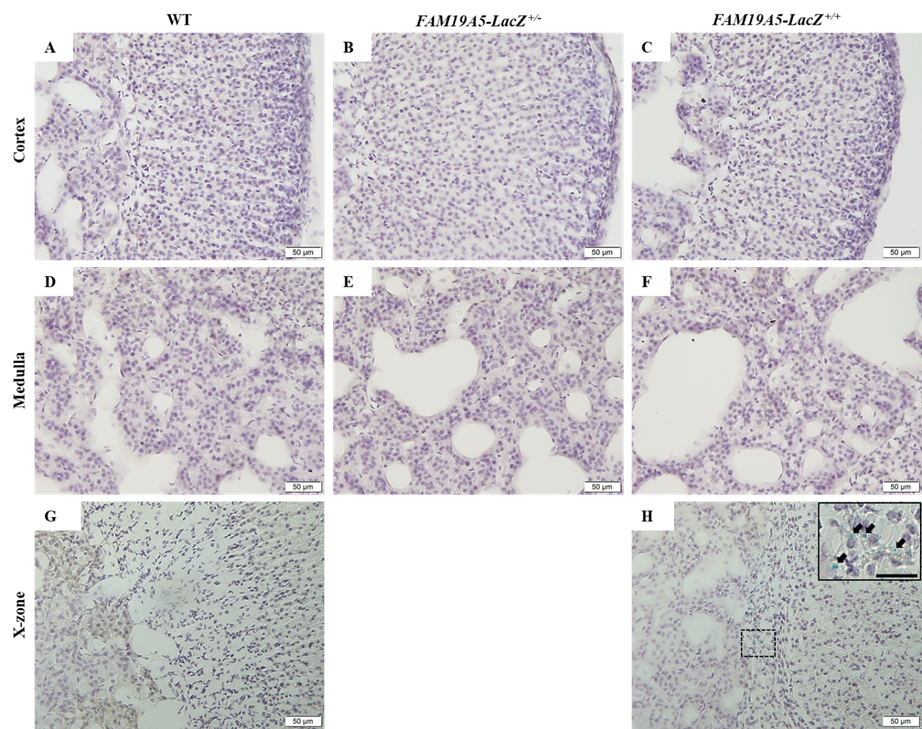

Representative light photomicrographs of adrenal gland cryosection of *(A, D, G)* wild type (WT, male #1 and female #1), *(B, E) FAM19A5-LacZ*^+/-^ (Heterozygote, male #1), and *(C, F, H)* *FAM19A5-LacZ*^+/+^ (Homozygote, male #1 and female #1). Adrenal cortex (A-C), medulla (D-F) and X-zone, located at the junction of the cortex and the medulla (G-H) were examined. C Cryosections were stained with X-gal solution and counter-stained with hematoxylin. No X-gal signals were determined in the adrenal cortex and medullar of WT and *FAM19A5-LacZ*^+/+^ mice. Punctate blue precipitates in X-Zone were observed in *FAM19A5-LacZ* KI mice but not in WT mice. Image in the dashed box is magnified in the inset with the same color as the dashed box. Black arrows indicate punctate blue precipitates. Scale bars in the inset represent 20 μm.

**Supplementary Fig. 9. Aorta X-gal signal.**

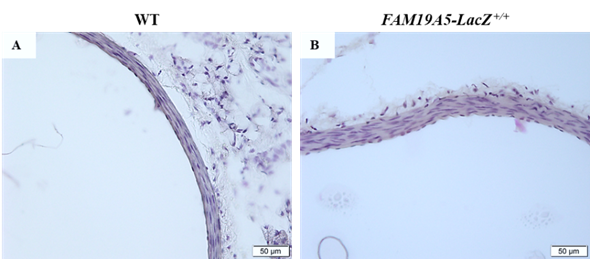

Representative light photomicrographs of thoracic (descending) aorta cryosection of *(A)* wild type (WT, male #2) and *(B) FAM19A5-LacZ*^+/+^ (Homozygote, male #2). Cryosections were stained with X-gal solution and counter-stained with hematoxylin. No X-gal signals were determined in both WT and *FAM19A5-LacZ*^+/+^ mice.

**Supplementary Fig. 10. Kidney X-gal signal.**

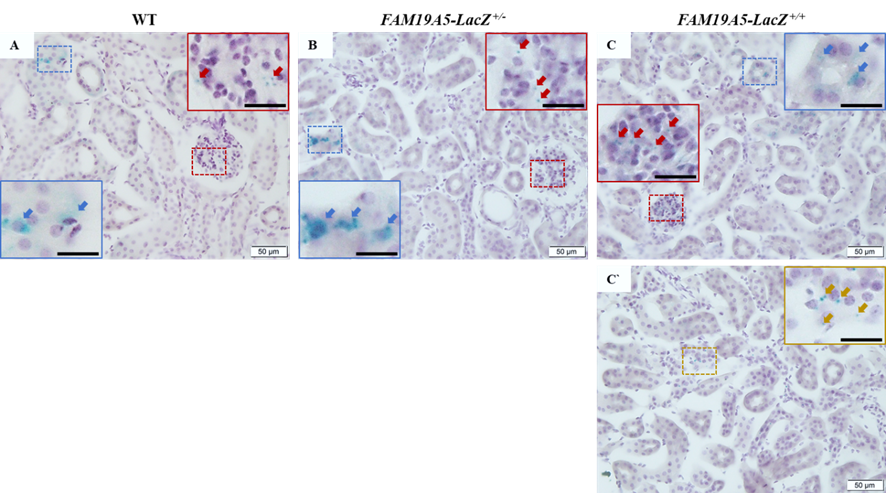

Representative light photomicrographs of kidney cryosection of *(A)* wild type (WT, male #2), *(B) FAM19A5-LacZ*^+/-^ (Heterozygote, male #1), and *(C* and *C`)* *FAM19A5-LacZ*^+/+^ (Homozygote, male #1). Cryosections were stained with X-gal solution and counter-stained with hematoxylin. Image in the dashed box is magnified in the inset with the same color as the dashed box. Red arrows indicate punctate blue precipitates in glomeruli observed in both WT and *FAM19A5-LacZ* KI mice. Blue arrows indicate dispersed blue precipitates in renal tubular epithelia observed in both WT and *FAM19A5-LacZ* KI mice. Yellow arrows indicate punctate blue precipitates in renal tubular epithelia observed only in *FAM19A5-LacZ* KI mice but not in WT mice. Scale bars in the inset represent 20 μm.

**Supplementary Fig. 11. Lung X-gal signal.**

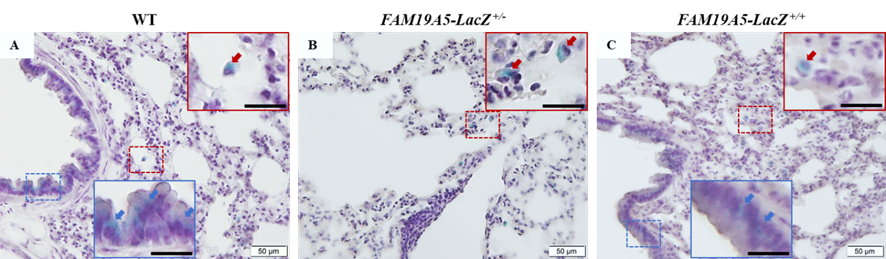

Representative light photomicrographs of lung cryosection of *(A)* wild type (WT, male #1), *(B)* *FAM19A5-LacZ*^+/-^ (Heterozygote, male #1), and *(C)* *FAM19A5-LacZ*^+/+^ (Homozygote, male #1). Cryosections were stained with X-gal solution and counter-stained with hematoxylin. Image in the dashed box is magnified in the inset with the same color as the dashed box. Red arrows indicate dispersed blue precipitates in alveolar macrophages observed in both WT and *FAM19A5-LacZ* mice. Blue arrows indicate dispersed blue precipitates in respiratory epithelia (pseudostratified columnar epithelia) observed in both WT and *FAM19A5-LacZ* KI mice. Scale bars in the inset represent 20 μm.

**Supplementary Fig. 12. Pancreas X-gal signal.**

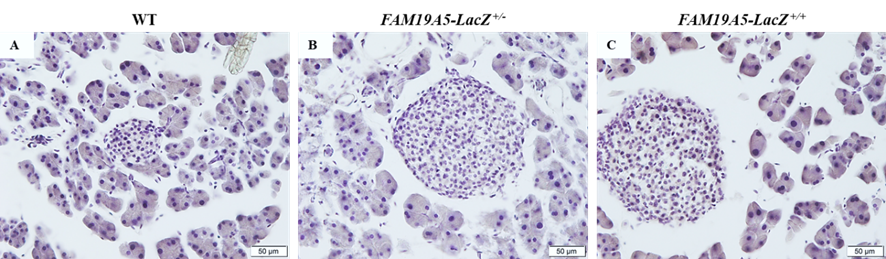

Representative light photomicrographs of pancreas cryosection of *(A)* wild type (WT, male #1), *(B) FAM19A5-LacZ*^+/-^ (Heterozygote, male #1), and *(C)* *FAM19A5-LacZ*^+/+^ (Homozygote, male #1). Cryosections were stained with X-gal solution and counter-stained with hematoxylin. No X-gal signals were determined in both WT and *FAM19A5-LacZ* KI mice.

**Supplementary Fig. 13. Liver X-gal signal.**

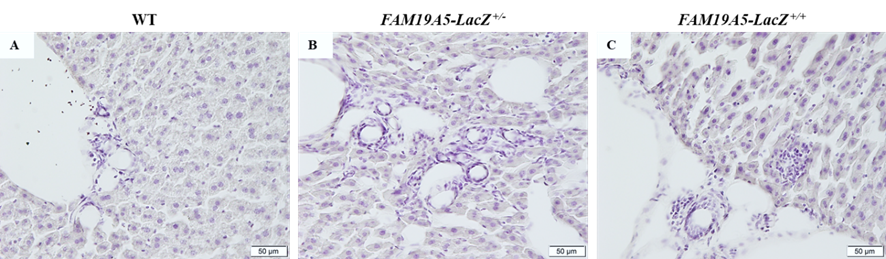

Representative light photomicrographs of liver cryosection of *(A)* wild type (WT, male #1), *(B) FAM19A5-LacZ*^+/-^ (Heterozygote, male #1), and *(C)* *FAM19A5-LacZ*^+/+^ (Homozygote, male #1). Cryosections were stained with X-gal solution and counter-stained with hematoxylin. No X-gal signals were determined in both WT and *FAM19A5-LacZ* KI mice.

**Supplementary Fig. 14. Salivary gland X-gal signal.**

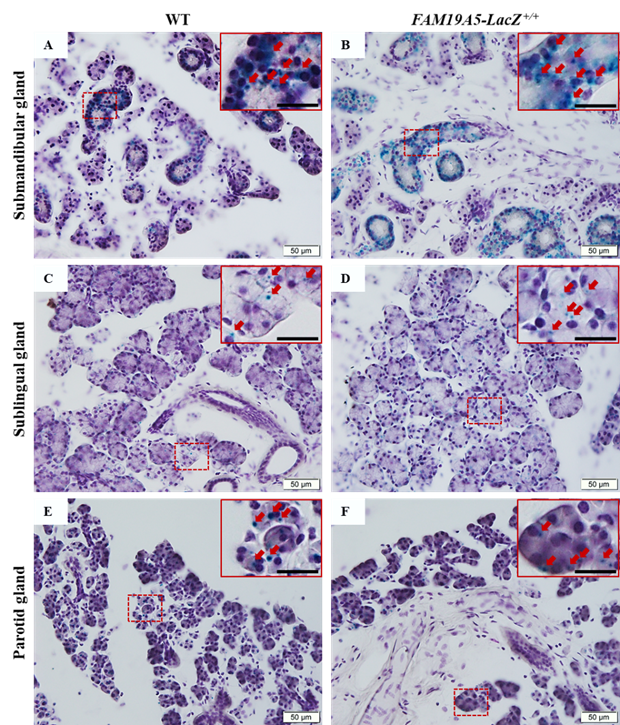

Representative light photomicrographs of salivary gland cryosection of *(A, C, E)* wild type (WT, male #1) and *(B, D, F)* *FAM19A5-LacZ*^+/+^ (Homozygote, male #2 and #3). Cryosections were stained with X-gal solution and counter-stained with hematoxylin. Image in the dashed box is magnified in the inset with the same color as the dashed box. *(A-B)* These images show submandibular gland of WT male #1 and *FAM19A5-LacZ*^+/+^ male #2. Red arrows indicate punctate/dispersed blue precipitates in mucous acini observed in both WT and *FAM19A5-LacZ*^+/+^ mice. *(C-D)* These images show sublingual gland of WT male #1 and *FAM19A5-LacZ*^+/+^ male #3. Red arrows indicate punctate blue precipitates in mucous acini observed in both WT and *FAM19A5-LacZ*^+/+^ mice. *(E-F)* These images show sublingual gland of WT male #1 and *FAM19A5-LacZ*^+/+^ male #2. Red arrows indicate punctate blue precipitates in serous acini observed in both WT and *FAM19A5-LacZ*^+/+^ mice. Scale bars in the inset represent 20 μm.

**Supplementary Fig. 15. Thymus X-gal signal.**

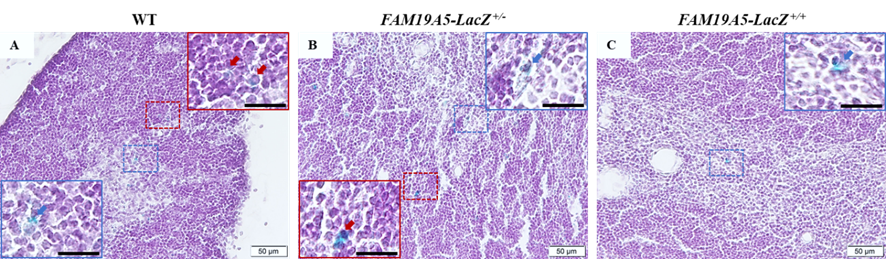

Representative light photomicrographs of thymus cryosection of *(A)* wild type (WT, male #1), *(B) FAM19A5-LacZ*^+/-^ (Heterozygote, male #1), and *(C)* *FAM19A5-LacZ*^+/+^ (Homozygote, male #1). Cryosections were stained with X-gal solution and counter-stained with hematoxylin. Image in the dashed box is magnified in the inset with the same color as the dashed box. Red arrows indicate dispersed blue precipitates in thymic cortex observed in both WT and *FAM19A5-LacZ* KI mice. Blue arrows indicate dispersed blue precipitates in thymic medulla observed in both WT and *FAM19A5-LacZ* KI mice. Scale bars in the inset represent 20 μm.

**Supplementary Fig. 16. Spleen X-gal signal.**

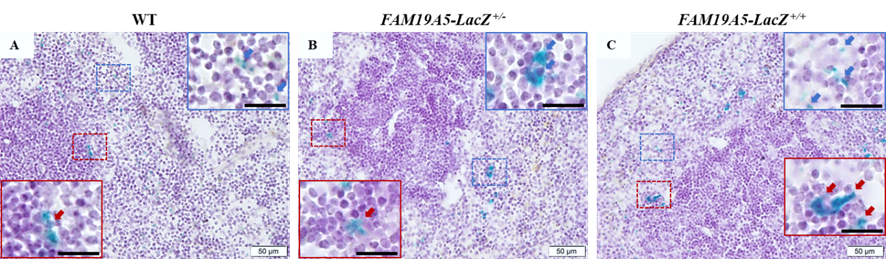

Representative light photomicrographs of spleen cryosection of *(A)* wild type (WT, male #1), *(B) FAM19A5-LacZ*^+/-^ (Heterozygote, male #1), and *(C)* *FAM19A5-LacZ*^+/+^ (Homozygote, male #1). Cryosections were stained with X-gal solution and counter-stained with hematoxylin. Image in the dashed box is magnified in the inset with the same color as the dashed box. Red arrows indicate dispersed blue precipitates in white pulp observed in both WT and *FAM19A5-LacZ* KI mice. Blue arrows indicate dispersed blue precipitates in red pulp observed in both WT and *FAM19A5-LacZ* KI mice. Scale bars in the inset represent 20 μm.

**Supplementary Fig. 17. Bone marrow X-gal signal.**

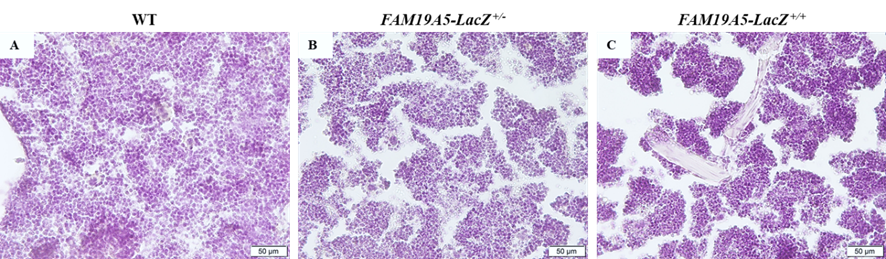

Representative light photomicrographs of femoral bone marrow cryosection of *(A)* wild type (WT, male #1), *(B) FAM19A5-LacZ*^+/-^ (Heterozygote, male #1), and *(C)* *FAM19A5-LacZ*^+/+^ (Homozygote, male #1). Cryosections were stained with X-gal solution and counter-stained with hematoxylin. No X-gal signals were determined in both WT and *FAM19A5-LacZ* KI mice.

**Supplementary Fig. 18. Pituitary gland X-gal signal.**

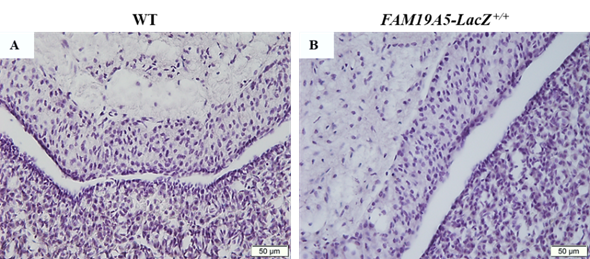

Representative light photomicrographs of pituitary gland cryosection of *(A)* wild type (WT, male #2) and *(B) FAM19A5-LacZ*^+/+^ (Homozygote, male #2). Cryosections were stained with X-gal solution and counter-stained with hematoxylin. No X-gal signals were determined in both WT and *FAM19A5-LacZ*^+/+^ mice.

**Supplementary Fig. 19. Thyroid gland X-gal signal.**

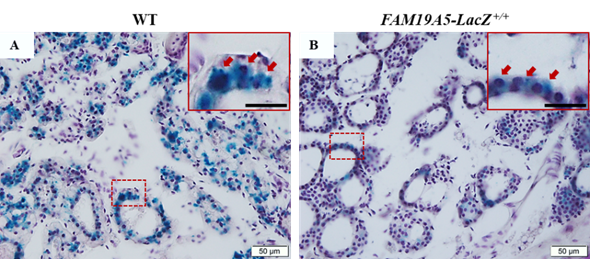

Representative light photomicrographs of thyroid gland cryosection of *(A)* wild type (WT, male #2) and *(B) FAM19A5-LacZ*^+/+^ (Homozygote, male #3). Cryosections were stained with X-gal solution and counter-stained with hematoxylin. Image in the dashed box is magnified in the inset with the same color as the dashed box. Red arrows indicate dispersed blue precipitates in follicular cells observed in both WT and *FAM19A5-LacZ*^+/+^ mice. Scale bars in the inset represent 20 μm.

**Supplementary Fig. 20. Skeletal muscle X-gal signal.**

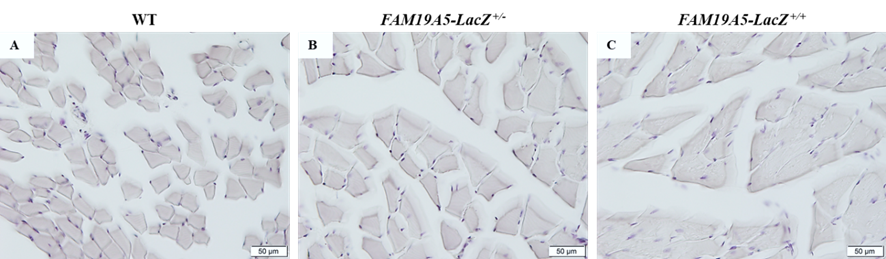

Representative light photomicrographs of femoral muscle cryosection of *(A)* wild type (WT, male #1), *(B) FAM19A5-LacZ*^+/-^ (Heterozygote, male #1), and *(C)* *FAM19A5-LacZ*^+/+^ (Homozygote, male #1). Cryosections were stained with X-gal solution and counter-stained with hematoxylin. No X-gal signals were determined in both WT and *FAM19A5-LacZ* KI mice.

**Supplementary Fig. 21. White adipose tissue X-gal signal.**

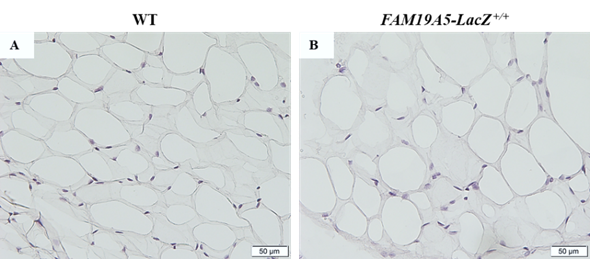

Representative light photomicrographs of white adipose tissue cryosection of *(A)* wild type (WT, male #1) and *(B) FAM19A5-LacZ*^+/+^ (Homozygote, male #1). All white adipose tissues were collected from the region adjacent to reproductive system, epididymis or ovary. Cryosections were stained with X-gal solution and counter-stained with hematoxylin. No X-gal signals were determined in both WT and *FAM19A5-LacZ*^+/+^ mice.

**Supplementary Fig. 22. Brown adipose tissue X-gal signal.**

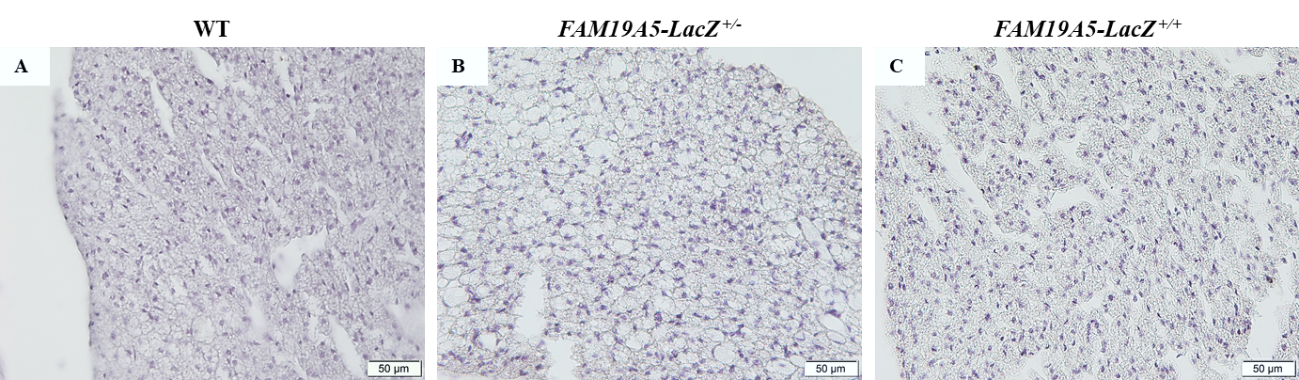

Representative light photomicrographs of brown adipose tissue cryosection of *(A)* wild type (WT, male #1), *(B) FAM19A5-LacZ*^+/-^ (Heterozygote, male #1), and *(C)* *FAM19A5-LacZ*^+/+^ (Homozygote, male #1). All brown adipose tissues were collected from the region adjacent to dorsal skin. Cryosections were stained with X-gal solution and counter-stained with hematoxylin. No X-gal signals were determined in both WT and *FAM19A5-LacZ* KI mice.

**Supplementary Fig. 23. Skin X-gal signal.**

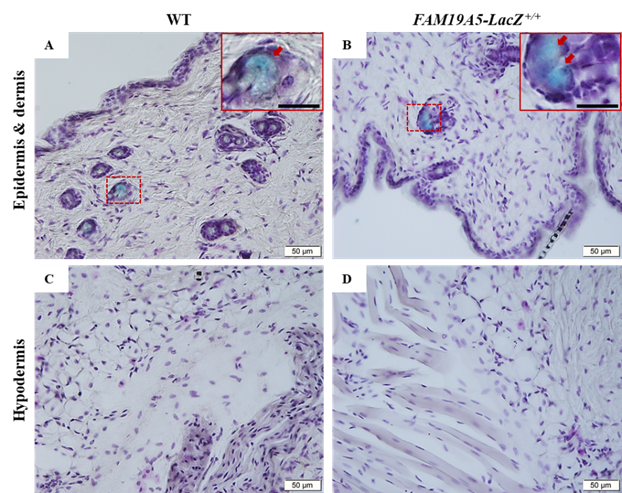

Representative light photomicrographs of skin cryosection of *(A, C)* wild type (WT, male #2) and *(B, D) FAM19A5-LacZ*^+/+^ (Homozygote, male #3). All skins were collected from the region of parietal/occipital bone. Cryosections were stained with X-gal solution and counter-stained with hematoxylin. *(A-B)* These images show epidermis and dermis regions of the skin. Image in the dashed box is magnified in the inset with the same color as the dashed box. Red arrows indicate dispersed blue precipitates in sebaceous glands observed in both WT and *FAM19A5-LacZ*^+/+^ mice. *(C-D)* These images show hypodermis regions of the skin. No X-gal signals were determined in both WT and *FAM19A5-LacZ*^+/+^ mice. Scale bars in the inset represent 20 μm.

**Supplementary Fig. 24. Seminal vesicle and coagulating gland X-gal signal.**

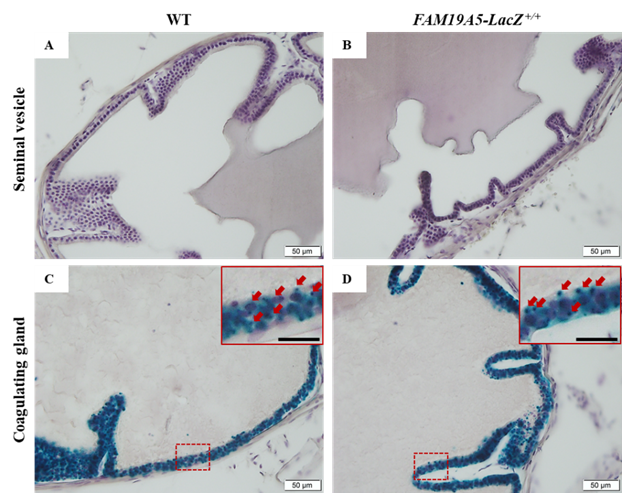

Representative light photomicrographs of seminal vesicle or coagulating gland cryosection of *(A, C)* wild type (WT, male #2) and *(B, D) FAM19A5-LacZ*^+/+^ (Homozygote, male #3). Cryosections were stained with X-gal solution and counter-stained with hematoxylin. *(A-B)* These images show seminal vesicles. No X-gal signals were determined in both WT and *FAM19A5-LacZ*^+/+^ mice. *(C-D)* These images show coagulating glands. Image in the dashed box is magnified in the inset with the same color as the dashed box. Red arrows indicate punctate blue precipitates in coagulating gland epithelia observed in both WT and *FAM19A5-LacZ*^+/+^ mice. Scale bars in the inset represent 20 μm.
